# CRISPR screens in physiologic medium reveal conditionally essential genes in human cells

**DOI:** 10.1101/2020.08.31.275107

**Authors:** Nicholas J. Rossiter, Kimberly S. Huggler, Charles H. Adelmann, Heather R. Keys, Ross W. Soens, David M. Sabatini, Jason R. Cantor

## Abstract

Forward genetic screens across hundreds of diverse cancer cell lines have started to define the genetic dependencies of proliferating human cells and how these vary by genotype and lineage. Most screens, however, have been carried out in culture media that poorly resemble metabolite availability in human blood. To explore how medium composition influences gene essentiality, we performed CRISPR-based screens of human cancer cell lines cultured in traditional versus human plasma-like medium (HPLM). Sets of medium-dependent fitness genes span several cellular processes and can vary with both natural cell-intrinsic diversity and the specific combination of basal and serum components that comprise typical culture media. Notably, we traced the causes for each of three conditional growth phenotypes to the availability of metabolites uniquely defined in HPLM versus traditional media. Our findings reveal the profound impact of medium composition on gene essentiality in human cells, and also suggest general strategies for using genetic screens in HPLM to uncover new cancer vulnerabilities and gene-nutrient interactions.

## INTRODUCTION

Loss-of-function forward genetic screens have been used to characterize protein function, map gene interaction networks, and define regulators of either drug or toxin resistance (Birsoy et al., 2015; Gilbert et al., 2014; Han et al., 2017; Kanarek et al., 2018; Kory et al., 2018; Shalem et al., 2013; Wang et al., 2014, 2017). There is also significant interest in leveraging such screens to identify the genetic dependencies of proliferating human cells, as these may suggest possible targets for cancer treatment (Tsherniak et al., 2017). Nonetheless, it is appreciated that genetic contributions to cell fitness may be context-dependent, and further, that gene essentiality is a quantitative property (Larrimore and Rancati, 2019; Rancati et al., 2017). Pooled loss-of-function screens based on either RNAi or CRISPR have been used in human cancer cell lines not only to establish a set of core essential genes, but also to define genetic dependencies that instead vary with genotype or cell lineage (Behan et al., 2019; Cheung et al., 2011; Hart et al., 2015; McDonald et al., 2017; Meyers et al., 2017; Tzelepis et al., 2016; Wang et al., 2015).

Environmental factors contribute to cell physiology and can also influence drug efficacy (Bader et al., 2020; Faubert et al., 2020; Luengo et al., 2017; Lyssiotis and Kimmelman, 2017; Muir and Vander Heiden, 2018). Recent *in vitro* studies have indeed demonstrated that cell fitness genes can vary with oxygen tension or culture in 3D spheroids versus 2D monolayers (Han et al., 2020; Jain et al., 2020). However, there has instead been little investigation into how the nutrient composition of cell culture media affects gene essentiality. Moreover, *in vitro* screens of human cells have been performed in traditional media that poorly recapitulate metabolite availability in human blood. (Ackermann and Tardito, 2019; Cantor, 2019).

Notably, progress has been made in conducting *in vivo* CRISPR-based screens, but such approaches also have limitations. Existing murine models recapitulate aspects of tumorigenesis and provide certain environmental factors that are often more difficult to model in standard culture systems, but genetic screens in mice are limited by cost, time, throughput, and control (Chow and Chen, 2018; Winters et al., 2018). There are also a number of differences in plasma metabolite levels between mice and humans, which could impact the physiology of human cells growing in mice (Cantor et al., 2017).

Previously, we developed a new culture medium (human plasma-like medium; HPLM) that contains over 60 polar metabolites and salt ions at concentrations that represent average values in adult human plasma (Cantor et al., 2017). Studies in human cancer cell lines and normal human T cells have demonstrated that HPLM has widespread effects on metabolism and other cellular processes (Cantor et al., 2017; Leney-Greene et al., 2020). Thus, we reasoned that by performing forward genetic screens in HPLM versus traditional media, it should be possible to identify genes differentially required for cells growing in biochemical conditions with greater relevance to human blood. This conditional essentiality paradigm has been illustrated in various microorganisms, as certain genes become critical for growth only in media that represent different laboratory or natural environments (Hillenmeyer et al., 2008; Nichols et al., 2010; Qian et al., 2012; Sassetti et al., 2001). By exploiting medium-dependent gene essentiality in human cells, it may be possible to develop strategies for treating cancer based on coupling targeted therapies with either dietary or enzyme-catalyzed modulation of plasma metabolites as well.

Here we perform CRISPR/Cas9 loss-of-function screens to systematically investigate how medium composition affects gene essentiality in human blood cancer cell lines. Analysis of these data reveals that sets of medium-dependent fitness genes encompass several cellular processes and can vary with the natural cell-intrinsic diversity of cancer cells, as well as with the combination of synthetic and serum components that comprise typical culture media. Follow-up work traces the causes of conditional growth phenotypes for each of glutamic-pyruvic transaminase 2 (GPT2), the mitochondrial pyruvate carrier (MPC), and glutaminase (GLS) to the availability of metabolites uniquely defined in HPLM versus standard media. Notably, the new gene-nutrient interaction identified for GLS is cell-specific and not completely predictive of *GLS* essentiality across different media conditions. By applying strategies that we describe, it should be possible to use CRISPR screens in HPLM to identify new targetable vulnerabilities, gene-nutrient interactions, and genetic drivers of other phenotypes (e.g. drug efficacy) in human cancers. Lastly, the causes responsible for medium-dependent essentiality across most hit genes we identify are unknown and of potential interest for future study.

## RESULTS AND DISCUSSION

### Genome-wide CRISPR/Cas9 screens reveal hundreds of medium-dependent fitness genes

Forward genetic screens based on either RNAi or CRISPR have been leveraged to define a set of core essential genes (CEGs) in human cancer lines and to catalog genetic dependencies that instead vary with natural cell-intrinsic diversity (Figure 1A) (Hart et al., 2015; McDonald et al., 2017; Tsherniak et al., 2017; Tzelepis et al., 2016; Wang et al., 2015, 2017). However, although environmental factors influence both cell physiology and gene essentiality, *in vitro* genetic screens have relied on culture media that most often consist of a basal medium with little relevance to the metabolite composition of human blood, and a fetal bovine serum (FBS) supplement that further contributes a typically undefined cocktail of additional components (Cantor, 2019). This point is well illustrated by the media conditions used across over 800 CRISPR screens from the DepMap project, of which over 75% were performed with one of two standard basal media (RPMI 1640, DMEM), and more than 80% carried out in media that contained a 10% FBS supplement (Figure 1B) (Dempster et al., 2019; Meyers et al., 2017).

**Figure 1.**
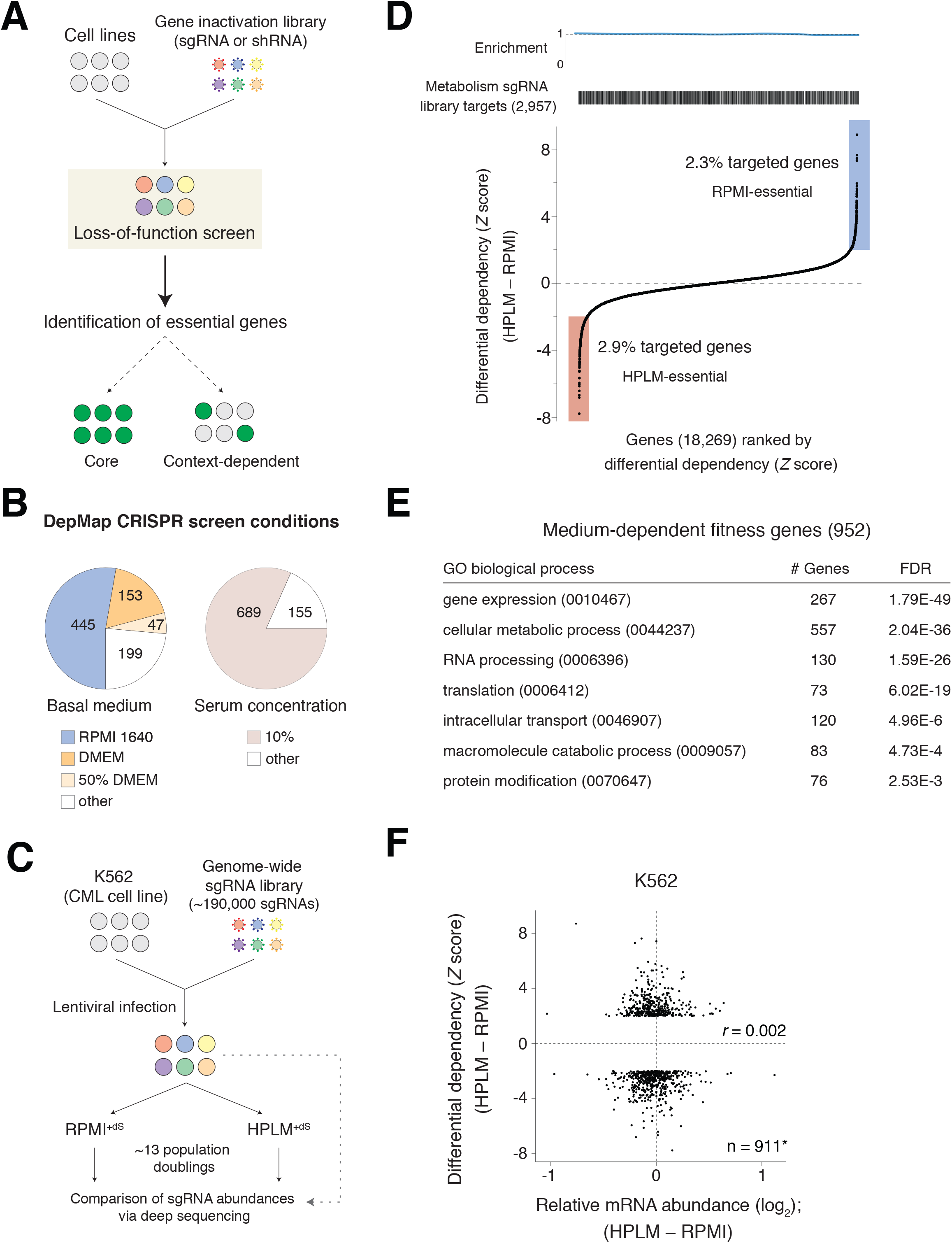
Genome-wide CRISPR screens for medium-dependent fitness genes. **See also Figure S1; Tables S1 and S2.** (A) Schematic for loss-of-function genetic screening methods applied to identify core or context-dependent essential genes. Gene inactivation is achieved by using either RNAi-based shRNA or CRISPR-based sgRNA libraries. sgRNA, single guide RNA. shRNA, short hairpin RNA. (B) Pie charts depicting culture compositions used across 844 CRISPR screens from DepMap. Fraction of screens performed in media containing RPMI 1640, DMEM, or an alternative basal medium (left). Fraction of screens performed in media containing 10% fetal bovine serum (FBS) (right). 50% DMEM refers to basal media that contained DMEM and another synthetic medium in a 1:1 mixture. (C) Schematic for CRISPR-based conditional essentiality profiling in K562 cells. RPMI^+dS^: RPMI 1640 with 5 mM glucose and 10% dialyzed FBS. HPLM^+dS^: HPLM with 10% dialyzed FBS. CML, chronic myeloid leukemia. (D) Plot of all targeted genes ranked by differential dependency score (see Methods). Barcode plot depicts the distribution of genes targeted by a metabolism-focused sgRNA library reported elsewhere (top). (E) Subset of enriched Gene Ontology (GO) pathways for the 952 medium-dependent hit genes analyzed using a PANTHER overrepresentation test (see also Table S2). (F) Relative expression versus differential dependency for medium-dependent hit genes. Values on the *x*-axis are relative mRNA abundance. *41 of the 952 hits were not identified in the RNA sequencing analysis. *r*, Pearson’s correlation coefficient.

We previously developed HPLM, a physiologic medium designed to more closely reflect the metabolic composition of human plasma (Cantor et al., 2017). To establish a complete HPLM-based medium, we add a 10% dialyzed FBS supplement (HPLM^+dS^) that provides various growth factors, hormones, and trace elements required to support cell proliferation, but minimizes the contribution of polar metabolites at otherwise undefined levels. Since RPMI 1640 (herein RPMI) has historically been the medium of choice for culturing human blood cells, we also created two RPMI-based reference media that each contain physiologic glucose (5 mM) but are supplemented with either 10% untreated FBS (RPMI^+S^) or 10% dialyzed FBS (RPMI^+dS^).

To test the hypothesis that proliferating human cells harbor medium-dependent essential genes, we used a genome-wide single guide(sg)RNA library (Wang et al., 2017) to perform CRISPR/Cas9 negative selection screens in the K562 chronic myeloid leukemia (CML) cell line. Following lentiviral infection and antibiotic selection in RPMI^+S^, cells were split and then passaged in either RPMI^+dS^ or HPLM^+dS^ – thus ensuring that causes for conditional growth phenotypes be restricted to differences in defined medium components (Figure 1C). Screens were passaged at the same frequency and cells doubled at near indistinguishable rates between the two conditions (Table S1). For each gene, we first calculated a gene score as the average log_2_-fold change in the abundance of all sgRNAs targeting the gene after 13 population doublings. By defining the median scores for sets of nontargeting sgRNAs and CEGs as 0 and −1, respectively, we then scaled all gene scores (Figure S1A) (Hart et al., 2017). Of note, the datasets from each screen could effectively discriminate CEGs from a distinct reference set of nonessential genes (Figure S1B) (Hart et al., 2014). By calculating a probability of dependency for each gene, we also found that the two datasets contained a comparable number of essential genes (probability > 0.5), which as anticipated, were enriched for roles in several fundamental cellular processes (Figure S1C) (Dempster et al., 1977, 2019).

We then standardized differential gene scores between the two conditions. By setting a *Z*-score cutoff of 2, we identified 427 RPMI-essential (positive) and 525 HPLM-essential (negative) genes, which collectively, were not enriched for targets of a metabolism-focused sgRNA library reported elsewhere (Figure 1D) (Birsoy et al., 2015). Pathway-enrichment analysis instead revealed that genes with strong conditional phenotypes represented many Gene Ontology (GO) biological processes, including metabolism, gene expression, RNA processing, and translation, among others (Figure 1E and Table S2). To ask whether potential differences in gene expression induced by HPLM^+dS^ versus RPMI^+dS^ could explain the growth phenotypes for candidate hits, we compared RNA sequencing from K562 cells following culture in each medium. However, we found no correlation between relative mRNA levels and differential dependency among the identified hit genes, indicating that a common proxy used to differentiate CEGs could not be similarly applied to identify medium-dependent fitness genes (Figure 1F and Table S2) (Wang et al., 2015).

### Medium-dependent fitness genes are involved in several cellular processes

Next, we designed a focused sgRNA library targeting 394 candidate hit genes (212 HPLM-essential and 182 RPMI-essential) and an additional 257 hit-related genes (e.g. shared pathway or family), and used it to similarly profile four blood cancer cell lines (K562, NOMO1, MOLM13, and SUDHL4) in HPLM^+dS^ versus RPMI^+dS^ (Figures 2A and S2A; Tables S3 and S4). Passaging frequencies were identical to those used for genome-wide screens and population doubling rates were comparable between the two conditions (Table S3). Since our focused sgRNA library contains a subset of nontargeting sgRNAs and others that target a fraction of the reference CEGs, we could analogously scale all secondary screen gene scores (Table S3). Importantly, we also confirmed that there was little influence on our genome-wide screen results if gene scores were instead scaled based on these two smaller sgRNA subsets. (Figure S2B and Table S1). Replicate secondary screen results in K562 cells were well correlated, and among those across all four cell lines, showed the highest correlation with genome-wide screens in the same medium (Figure S2C and S2D). Therefore, we combined data from the two replicates to establish pooled datasets and corresponding differential dependencies that were highly correlated with those from our genome-wide screen results, indicating that conditional growth phenotypes could be largely recapitulated by screening with the focused sgRNA library (Figures 2B and S2E).

**Figure 2.**
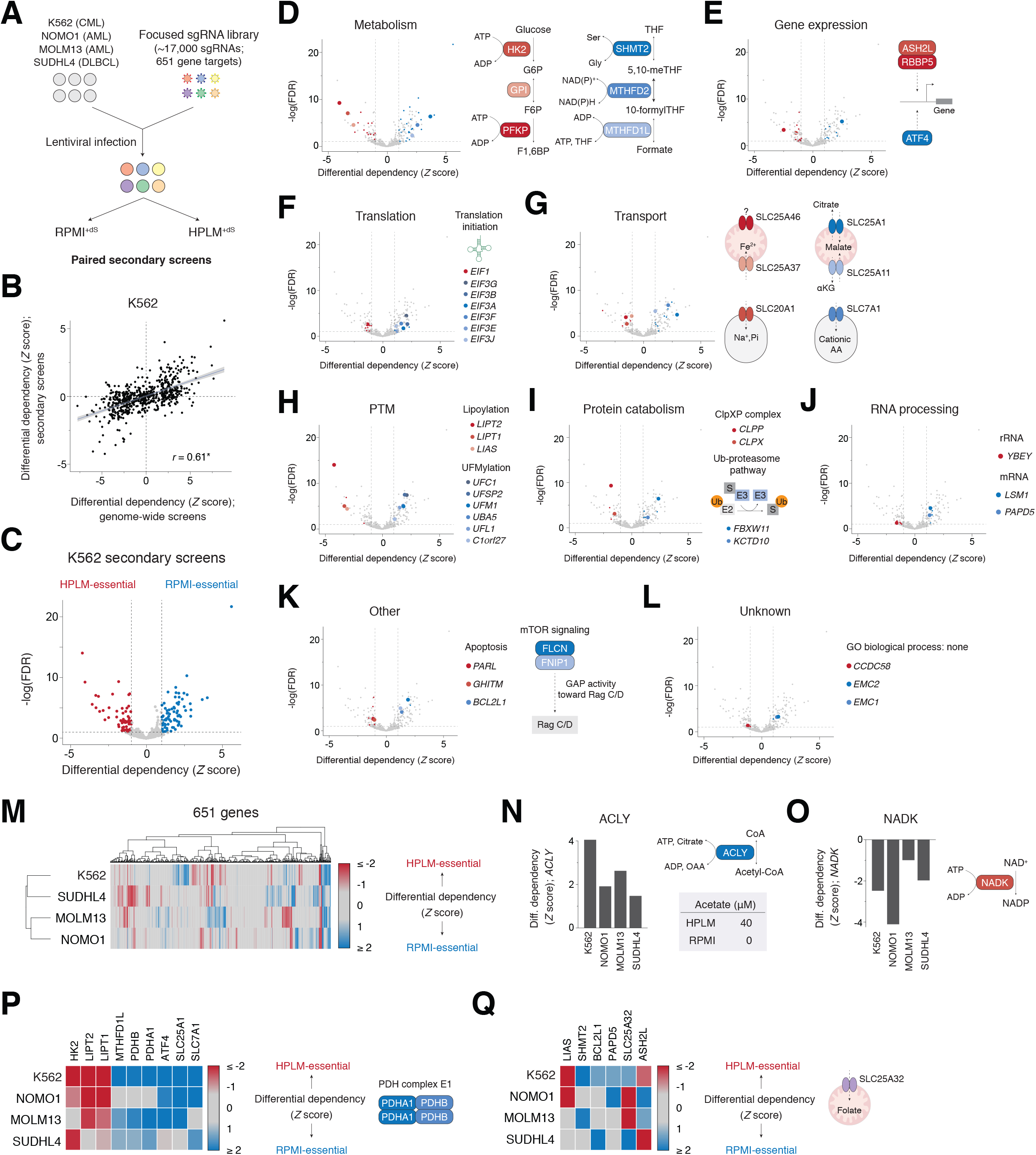
Medium-dependent fitness genes are involved in several cellular processes and can vary with natural cell-intrinsic diversity. **See also Figures S2 and S3; Tables S3 and S4** (A) Schematic for secondary screens in each of four human blood cancer lines using a focused sgRNA library. AML, acute myeloid leukemia; DLBCL, diffuse large B-cell lymphoma. (B) Comparison of conditional phenotypes for 651 genes in genome-wide versus secondary K562 screens. Data are fit by linear regression (blue line); shaded bands indicate 95% confidence intervals. *r*, Pearson’s correlation coefficient. \**P* = 2.2 × 10^−16^. Secondary screen data for the K562 cell line are from pooled replicates in all panels. (C) HPLM-essential (red) and RPMI-essential (blue) genes identified from secondary K562 screen results. Dashed lines indicate standard scores of ± 1 (*x*-axis) and a false discovery rate (FDR) equivalent to 0.1 (*y*-axis). (D-L) Medium-dependent fitness genes in K562 cells encode proteins involved in (D) metabolism, (E) gene expression, (F) translation, (G) transport, (H) posttranslational modification (PTM), (I) protein catabolism, (J) RNA processing, and (K) other cellular processes, including apoptosis and mTOR signaling. Other hits are not currently annotated for any GO biological process (L). Shaded points in each panel represent genes manually curated for involvement in the cellular process. Enlarged points are further highlighted (right). (M) Cluster map showing conditional phenotypes for genes targeted by the focused sgRNA library across the four screened cell lines. (N-O) Conditional phenotypes for *ACLY* (N) and *NADK* (O) from secondary screen results (left). Reactions catalyzed by each encoded enzyme (right). Defined acetate levels in HPLM and RPMI are also indicated (N). (P) Heatmap of conditional phenotypes for the indicated genes (left). PDHA1 and PDHB are components of the PDH complex E1 subunit (right). Remaining genes are highlighted elsewhere in the Figure. (Q) Heatmap of conditional phenotypes for the indicated genes (left). SLC25A32 is mitochondrial folate transporter (right). Remaining genes are highlighted elsewhere in the Figure.

First, to identify hit genes in K562 cells that were most likely to be biologically meaningful, we tested the significance of each differential dependency from our pooled datasets. By setting a standard score cutoff of 1, we identified 78 RPMI-essential and 71 HPLM-essential hits at a 0.1 false discovery rate (FDR) (Figure 2C). Given the targeting bias of our focused library, this cutoff was chosen to maintain selection for differential gene scores that met the applied threshold used to define conditional essentiality in our genome-scale screen results. Consistent with the pathway enrichment analysis above, we found that genes with strong conditional phenotypes represented several cellular processes.

Despite normalized glucose availability between HPLM^+dS^ and RPMI^+dS^, metabolic genes encoding enzymes that catalyze initial steps of glycolysis (*HK2, GPI*, and *PFKP*) scored as HPLM-essential, while others that can catalyze successive reactions in one-carbon metabolism (*SHMT2, MTHFD2*, and *MTHFD1L*) instead scored as RPMI-essential – perhaps given the uniquely defined availability of formate in HPLM (Figure 2D). We also uncovered medium-dependent fitness genes associated with gene expression, including a reported heterodimer with histone methyltransferase activity *(ASH2L and RBBP5)* and a transcription factor involved in the cellular response to nutrient restriction *(ATF4)* (Figure 2E) (Cao et al., 2010). Interestingly, whereas the translation initiation factor 1 gene *(EIF1)* was identified as HPLM-essential, several others encoding members of the translation initiation factor 3 complex instead scored as RPMI-essential (Figure 2F).

The conditional essentiality analysis also revealed hits involved in the transport of various metabolites *(SLC25A1, SLC25A11*, and *SLC7A1)*, small ions *(SLC25A37* and *SLC20A1)*, or even an unknown substrate *(SLC25A46)* (Figure 2G). Genes involved in lipoylation, a posttranslational attachment of lipoamide to proteins, were identified as HPLM-essential hits, while others that encode components of the UFMylation machinery, a system that attaches UFM1 to proteins, instead scored as RPMI-essential (Figure 2H) (Komatsu et al., 2004; Rowland et al., 2018; Solmonson and DeBerardinis, 2018; Wang et al., 2017). Additional genes with strong medium-dependent phenotypes are involved in protein catabolism, including components of the ClpXP protease complex *(CLPX* and *CLPP)* and of E3 ubiquitin-protein ligase complexes *(KCTD10* and *FBXW11)*, as well as in RNA processing *(YBEY, LSM1*, and *PAPD5)*, apoptosis *(BCL2L1, PARL*, and *GHITM)*, and the mTOR pathway *(FLCN* and *FNIP1)* (Tsun et al., 2013), among other processes (Figures 2I, 2J, and 2K).

Lastly, we could also identify hit genes with no annotated function, such as *CCDC58* and multiple members of the endoplasmic reticulum membrane protein complex *(EMC1* and *EMC2)* (Figure 2L). While the causes underlying most conditional phenotypes identified from our analysis are not immediately apparent, these results do reveal that defined basal medium composition has a dramatic impact on gene essentiality in human cells.

### Medium-dependent gene essentiality can be influenced by cell-intrinsic factors

Next, we sought to ask how conditional gene essentiality is affected by natural cell-intrinsic diversity. Overall, the sets of genes with strong medium-dependent phenotypes showed variable overlap and a number of distinct patterns across our cell line panel (Figure 2M). We observed a positive growth phenotype in each cell line for *ACLY*, which encodes an enzyme that generates acetyl-CoA from citrate, likely given that HPLM contains defined acetate, a substrate that supports an alternative cellular route to acetyl-CoA synthesis (Figure 2N) (Zhao et al., 2016). In contrast, the cytosolic NAD kinase gene *(NADK)* was identified as a common HPLM-essential hit across all four cell lines, though the gene-nutrient interaction responsible for this effect is not immediately clear (Figure 2O).

We also uncovered several genes with conditional phenotypes shared among only three cell lines, including those that encode components of the pyruvate dehydrogenase complex E1 subunit *(PDHA* and *PDHB)* (Figure 2P). Further, four of the six genes that encode enzymes along the de novo purine biosynthesis pathway scored as RPMI-essential, likely owing to HPLM-specific availability of defined hypoxanthine, a salvage substrate for purine synthesis (Figure S3A). Differences in hypoxanthine uptake rates may explain why this pattern was not observed in K562 cells. Surprisingly, three genes involved in methionine salvage *(MTR, MTRR*, and *MMACHC)* were identified as positive hits in K562 cells, but instead scored as HPLM-essential in each of the three remaining cell lines, highlighting how cell-intrinsic heterogeneity can influence metabolic responses to differential metabolite availability (Figure S3B).

Our analysis between cell lines also revealed hit genes shared by only two cell lines, such as the mitochondrial folate transporter *(SLC25A32)* (Figure 2Q), and many others restricted to a single cell line and representing a number of processes (Figure S3C). Together, our results are consistent with the notion that gene essentiality depends on a combination of cell-intrinsic and environmental factors (Rancati et al., 2017), suggesting that CRISPR screens in HPLM across broader cell line panels should reveal new cancer-specific vulnerabilities.

### Identification of a gene-nutrient interaction between *GPT2* and alanine

While the overall sets of conditional fitness genes varied within our panel, we found that *GPT2* was the most strongly scoring RPMI-essential hit in each cell line (Figure 3A). Interestingly, however, *GPT2* is annotated as an essential gene in fewer than 1% of the nearly 800 CRISPR screens in DepMap (Figure S4A) (Dempster et al., 2019; Meyers et al., 2017). *GPT2* encodes one of two enzymes that catalyze the reversible conversion of pyruvate and glutamate to alanine and α-ketoglutarate (αKG), but that differ in their subcellular localization (GPT1, cytosolic; GPT2, mitochondrial) (Figure 3B). For each cell line, our secondary screen results suggested that loss of *GPT2* had little influence on cell growth in HPLM^+dS^ but caused markedly negative effects in RPMI^+dS^, whereas *GPT1* deletion had little influence in either condition (Figures S4B and S4C). Consistent with these latter results, *GPT1* is annotated as an essential gene in the same negligible fraction of DepMap CRISPR screens as *GPT2* (Figure S4D). Notably, RNA sequencing across nearly 1,300 human cancer cell lines indicate near absolute selective expression of *GPT2* versus *GPT1* (Figure S4E) (Ghandi et al., 2019). For the cell lines in our panel, we confirmed this relative expression phenotype at the protein level and also found that GPT2 abundance in K562 cells was unaffected by culture in HPLM^+dS^ relative to either RPMI-based medium (Figures 3C and 3D).

**Figure 3.**
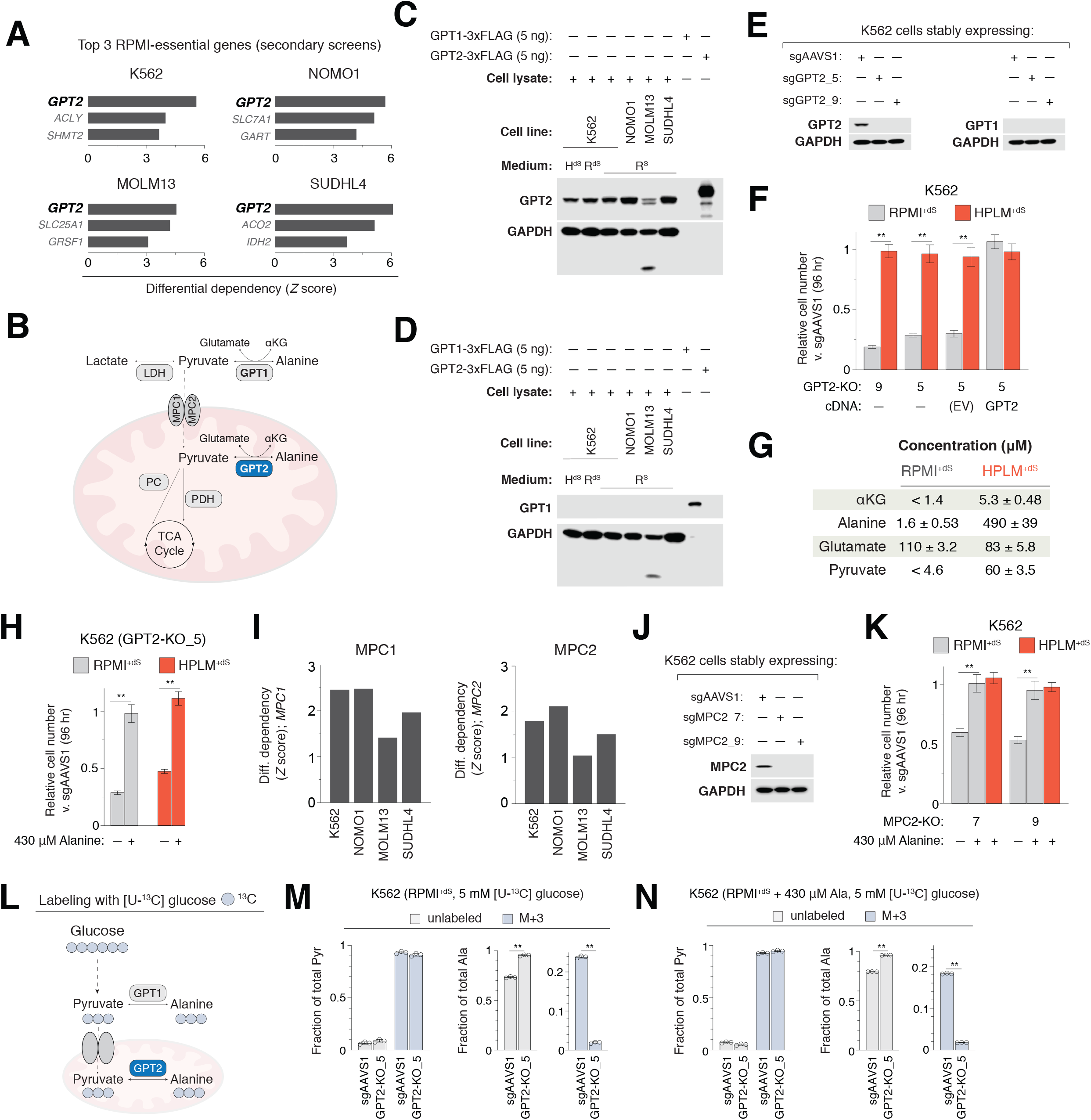
Identification of a gene-nutrient interaction between GPT2 and alanine. **See also Figure S4 and Table S5** (A) Top three most strongly scoring RPMI-essential hits from secondary screen results across four screened cell lines. Secondary screen data for the K562 cell line are from pooled replicates. (B) Schematic depicting various cellular fates for pyruvate and the reversible reactions catalyzed by GPT enzymes. The mitochondrial pyruvate carrier (MPC) is an obligate heterodimer (MPC1 and MPC2) that transports pyruvate into the mitochondrial matrix. αKG, α-ketoglutarate; GPT, glutamic-pyruvic transaminase; LDH, lactate dehydrogenase; PDH, pyruvate dehydrogenase; PC, pyruvate carboxylase. (C-D) Immunoblots for expression of GPT1 (C) and GPT2 (D). Lanes containing purified GPT1-3xFLAG and GPT2-3xFLAG confirm primary antibody specificity. GAPDH served as a loading control. Of note, we suspect that high intrinsic protease activity in the MOLM13 cell line causes the displayed banding depicted for both GPT2 and GAPDH. (E) Immunoblots for expression of either GPT2 (left) or GPT1 (right) in *GPT2*-knockout (sgGPT2) and control K562 cells transduced with sgAAVS1. GAPDH served as a loading control. (F) Relative growth of *GPT2*-knockout versus control cells in the indicated conditions (mean ± SD, *n* = 3, \*\**P* < 0.005). Knockout cells transduced with an expression construct either containing a sgRNA-resistant *GPT2* (GPT2) or lacking a cDNA (EV, empty vector). (G) Concentrations of GPT reaction components in each of RPMI^+dS^ and HPLM^+dS^ as measured by metabolite profiling (mean ± SD, *n* = 3). Of note, neither αKG nor pyruvate could be readily detected in RPMI^+dS^ with the applied method; indicated threshold concentrations correspond to levels measured in RPMI^+S^. (H) Relative growth of *GPT2*-knockout versus control cells in the indicated conditions (mean ± SD, *n* = 3, \*\**P* < 0.005). (I) Conditional phenotypes for *MPC1* (left) and *MPC2* (right) from secondary screen results. (J) Immunoblot for expression of MPC2 in both *MPC2*-knockout cells (sgMPC2) and control K562 cells. GAPDH served as a loading control. (K) Relative growth of *MPC2*-knockout versus control cells in the indicated conditions (mean ± SD, *n* = 3, \*\**P* < 0.005). (L) Schematic depicting the incorporation of ^13^C from glucose into alanine via pyruvate. (M-N) Fractional labeling of pyruvate (left) and alanine (right) following 24 hr culture of cells in RPMI^+dS^ containing [U-^13^C]-glucose (M) and further supplemented with 430 μM alanine (N) (mean ± SD, *n* = 3, \*\**P* < 0.005). M+3, incorporation of three ^13^C.

To examine the conditional phenotype for *GPT2* deletion, we engineered *GPT2*-knockout K562 clonal cells, which had the same lack of detectable GPT1 expression as control K562 cells transduced with an AAVS1-targeting sgRNA (Figure 3E) (Wang et al., 2015). By using short-term growth assays, we confirmed that *GPT2* deletion caused a pronounced growth defect specific to culture in RPMI^+dS^ (Figure 3F). Importantly, the expression of a sgRNA-resistant *GPT2* cDNA fully rescued this defect, while transduction with the same construct lacking a cDNA did not.

To determine why *GPT2* deletion impaired cell growth in RPMI^+dS^, we first considered the relative availability of each GPT reaction component, reasoning that de novo synthesis of one or more could become essential under limiting conditions. While glutamate levels between the two media are comparable, HPLM contains the three remaining components at concentrations at least 5-(αKG), 15-(pyruvate), and 200-fold (alanine) greater than those quantified in RPMI^+dS^ (Figure 3G). Studies in human cancer cells over the past decade have highlighted a role for GPT2 in facilitating glutamine anaplerosis via the production of αKG (Hao et al., 2016; Kim et al., 2019; Smith et al., 2016; Weinberg et al., 2010), and further, others have reported that GPT can serve to fuel the TCA cycle in certain cancers by catabolizing alanine to pyruvate (Parker et al., 2020; Sousa et al., 2016). Therefore, we considered whether the differential availability of either αKG or pyruvate could explain the RPMI-essential phenotype of *GPT2* deletion. Among the stocks of pooled HPLM components that we create is one containing αKG, pyruvate, and eight additional water-soluble acids (WSAs) (Figure S4F and Table S5). However, the addition of this WSAs pool to RPMI^+dS^ could not boost the relative growth of *GPT2*-knockout cells, and neither could that of cell-permeable dimethyl αKG (DM-αKG) at levels up to 40-fold greater than those of αKG in HPLM (Figure S4G). We then considered the relative availability of alanine which, despite being the second most abundant amino acid in human blood, is not a defined component of either RPMI or DMEM. When we supplemented RPMI^+dS^ with physiologic alanine (430 μM), we observed a full rescue of the growth defect and, in addition, also found that the removal of alanine from HPLM^+dS^ could impair the relative growth of *GPT2*-knockout cells as well (Figure 3H).

Next, we reasoned that GPT2-catalyzed alanine production would require that GPT2 have access to the corresponding reaction substrates. The mitochondrial pyruvate carrier (MPC) is an obligate heterodimer (MPC1 and MPC2) that transports pyruvate into the mitochondrial matrix (Bricker et al., 2012; Herzig et al., 2012). Consistent with our rationale, both *MPC1* and *MPC2* were identified as RPMI-essential hits in all four cell lines (Figure 3I). To confirm this conditional growth phenotype, we engineered *MPC2*-knockout K562 clonal cells, which as expected, showed a growth defect specific to culture in RPMI^+dS^ that could be similarly rescued by the addition of physiologic alanine (Figures 3J and 3K).

To examine the contribution of GPT2 to the intracellular alanine pool, we next compared ^13^C-labeling patterns of pyruvate and alanine following 24 hr culture of *GPT2*-knockout and control cells in RPMI^+dS^ containing [U-^13^C]-glucose (Figure 3L). *GPT2* deletion had effectively no influence on the fraction of pyruvate labeled with three ^13^C (M+3) but decreased that of M+3-alanine to near negligible levels (Figure 3M). These results are consistent with prior studies that report a reduced fractional isotopic labeling of alanine downstream of glucose/pyruvate in either *Gpt2*-null mouse embryonic fibroblasts or human cells treated with an MPC inhibitor (Ouyang et al., 2016; Vacanti et al., 2014; Yang et al., 2014). By instead performing our glucose tracing experiments in RPMI^+dS^ containing 430 μM alanine, we observed only minor effects on the M+3-labeling patterns above, indicating that the GPT2-mediated production of alanine and αKG is not necessarily dictated by alanine availability (Figure 3N).

Together, these results reveal that medium-dependent growth phenotypes for *GPT2* and *MPC1/2* could be traced to the differential availability of alanine, one of the three GPT reaction components uniquely defined in HPLM versus RPMI. By comparing gene essentiality profiles in HPLM^+dS^ and RPMI^+dS^, we found that both GPT2 and the MPC serve important roles in supporting an alanine-dependent cell-essential demand in conditions of relative alanine limitation.

### GPT2 supports protein synthesis under conditions of alanine restriction

To begin to determine the cell-essential demand supported by de novo alanine production for cells growing in RPMI^+dS^, we performed unbiased metabolite profiling in *GPT2*-knockout and control cells following 24 hr culture in either HPLM^+dS^ or RPMI^+dS^ (Table S6). Of note, among the three GPT reaction components uniquely provided in HPLM, only the relative levels of measured alanine in the control cells reflected availability differences between the two media (Figure S5A). Interestingly, *GPT2* deletion had widespread effects on cellular metabolite abundances following culture in RPMI^+dS^ but not in HPLM^+dS^ (Figure 4A).

**Figure 4.**
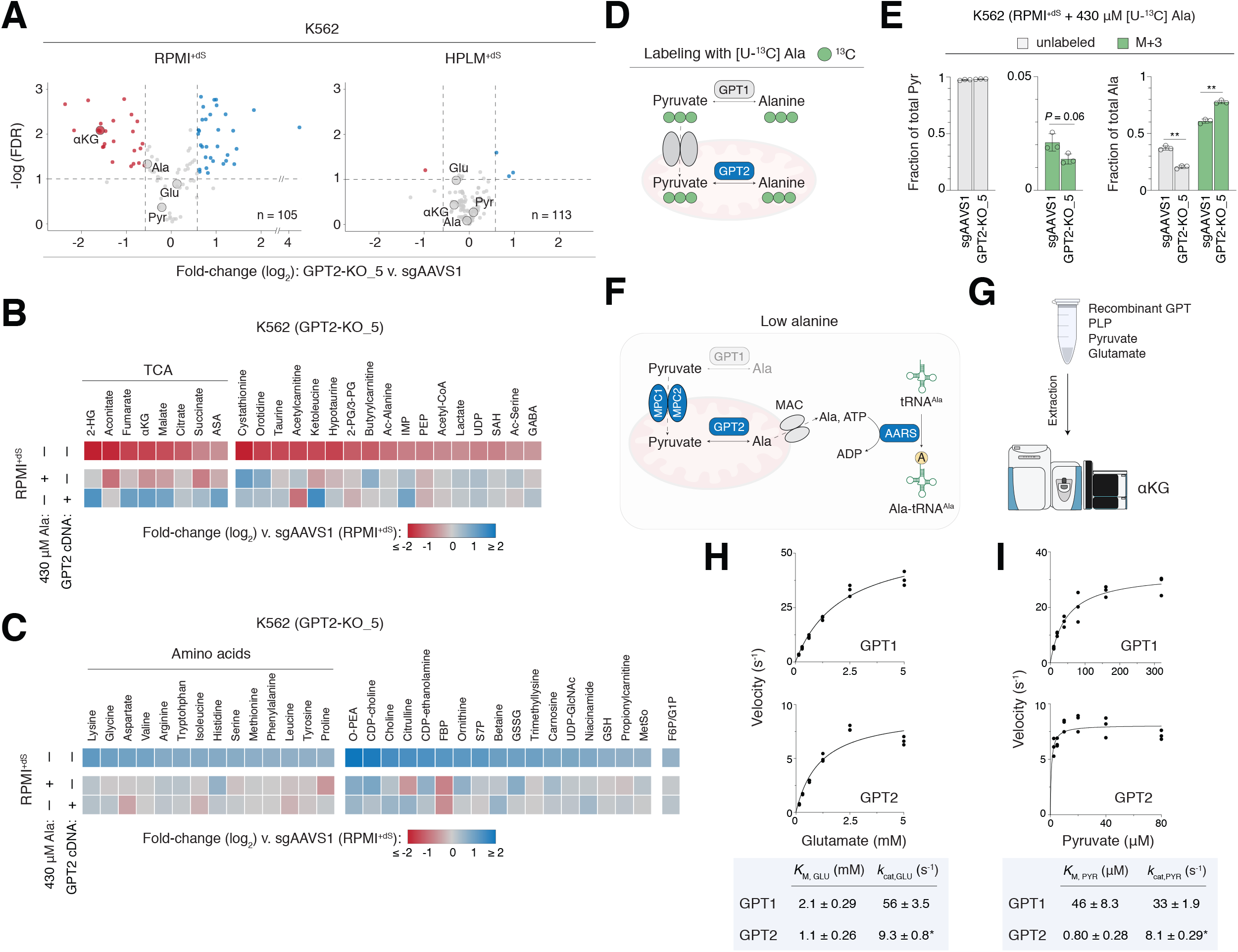
GPT2 supports protein synthesis under conditions of alanine restriction and human GPTs exhibit markedly different *K*_M_ values for pyruvate. **See also Figures S5 and S6; Table S6** (A) Volcano plots of cellular metabolite abundances in *GPT2*-knockout versus control K562 cells following culture in either RPMI^+dS^ (left) or HPLM^+dS^ (right) (*n* = 3). Each point represents one metabolite. Dashed lines indicate fold-changes equivalent to ± 1.5 (*x*-axis) and FDR equivalent to 0.1 (*y*-axis). GPT reaction components are enlarged and labeled. (B-C) Heatmap of relative cellular abundances for metabolites highlighted in either red (B) or blue (C) following culture in RPMI^+dS^ in panel A. Knockout cells following culture in each of the indicated conditions (top and middle rows) or transduced with *GPT2* cDNA (bottom row) versus control cells in RPMI^+dS^. Metabolite groups in each panel are sorted by log_2_-transformed fold change of the top row. Argininosuccinic acid (ASA) is a precursor to fumarate. Remaining metabolite abbreviations in Table S6. (D) Schematic depicting the incorporation of ^13^C from alanine into pyruvate. (E) Fractional labeling of pyruvate (left) and alanine (right) following 24 hr culture of cells in RPMI^+dS^ containing 430 μM [U-^13^C]-alanine (mean ± SD, *n* = 3, \*\**P* < 0.005). M+3, incorporation of three ^13^C. (F) Schematic depicting the proposed model for the cell-essential role of GPT2 in conditions of relative alanine limitation. GPT2 generates de novo alanine that exits mitochondria and charges the cytosolic alanyl-tRNA synthetase (AARS) to support translation. Proteins encoded by RPMI-essential hit genes (blue). A canonical mitochondrial alanine carrier (MAC) has not yet been identified, though the reported serine transporter SFXN1 shows promiscuous transport activity for alanine (Kory et al., 2018). (G) Schematic depicting an in vitro GPT assay in the direction of alanine and αKG formation using LC-MS-based detection of αKG. (H-I) Plots of reaction velocity as a function of either glutamate (H) or pyruvate (I) concentration for human GPT1 (top) and GPT2 (right) (*n* = 3). Data are fit by Michaelis-Menten curves to estimate *K*_M_ and *k*_cat_ for each substrate (bottom). *measured *k*_cat_ values for GPT2 may be underestimated (see Main Text).

By setting a fold-change cutoff of −1.5 and 0.1 FDR, we found that *GPT2* knockout reduced the levels of 25 metabolites, including αKG and several others associated with the TCA cycle but not alanine, whose abundance was instead reduced by an extent half that of αKG. However, while the expression of our *GPT2* cDNA reversed most of the αKG-related changes, culture in RPMI^+dS^ supplemented with physiologic alanine did not (Figure 4B). We also observed that *GPT2* knockout increased the abundances of 32 metabolites by at least 1.5-fold at the same FDR, among which nearly half were proteinogenic amino acids. Notably, these changes were instead largely rescued by both *GPT2* cDNA expression and alanine supplementation, suggesting they were more likely relevant to the gene-nutrient interaction between *GPT2* and alanine (Figures 4C and S5B). Given that reduced translation and increased cellular amino acid pools are each primary consequences of amino acid restriction (Bröer and Bröer, 2017), these results suggest that *GPT2*-knockout cells growing in RPMI^+dS^ exhibit a metabolic phenotype consistent with nutrient restriction.

Next, to ask if GPT2 might support a cell-essential catabolic demand for cells growing in conditions of relative alanine restriction, we compared the ^13^C-labeling of alanine and pyruvate following 24 hr culture of *GPT2*-knockout and control K562 cells in RPMI^+dS^ containing 430 μM [U-^13^C]-alanine (Figure 4D). Differences in fractional M+3-alanine labeling complemented those from our glucose tracing results as expected, but *GPT2* deletion had little impact on the otherwise minimal fraction of M+3-pyruvate measured in our control cells (Figure 4E). Fractional labeling of additional metabolites downstream of pyruvate metabolism were near negligible as well (Table S6). Of note, the addition of physiologic alanine could also fully rescue the relative growth defect of *GPT2*-knockout cells in RPMI^+dS^ following 24 hr culture, importantly indicating that the unbiased metabolite profiling and tracing results analyzed at this timepoint were biologically relevant to the gene-nutrient interaction (Figure S5C).

Collectively, these results suggest a model in which alanine supports the non-catabolic cell-essential demand of protein synthesis, a role similarly proposed in the context of normal T-cell activation (Figure 4F) (Ron-Harel et al., 2019). Consistent with this model, *GPT2* and the alanyl-tRNA synthetase gene *(AARS)* scored as the top and third most strongly scoring RPMI-essential hits, respectively, from our genome-wide screen results in K562 cells (Figure S5D). This conditional phenotype for loss of *AARS* was recapitulated in secondary screes and observed for two additional cell lines as well (NOMO1, MOLM13) (Figure S5E). Notably, *AARS* is among the reference CEGs (Hart et al., 2017), and indeed, its loss caused a marked growth defect in each screen condition (Figure S5F). To reconcile these data, we speculate that from the point of Cas9-mediated *AARS* cleavage, cellular alanine concentrations above a critical threshold can maintain tRNA^Ala^ charging until the turnover of residual AARS at rates that likely vary between cell lines. Interestingly, the related alanyl-tRNA synthetase involved in mitochondrial translation *(AARS2)* did not similarly score as a strong medium-dependent fitness gene.

### Human GPTs exhibit markedly different *K*_M_ values for pyruvate

Next, given the relative expression of *GPT2* versus *GPT1* across most human cancer cell lines, we considered if enforced expression of *GPT1* might complement *GPT2* deletion. When we transduced *GPT2*-knockout cells with a *GPT1* cDNA, we observed complete rescue of the growth defect in RPMI^+dS^, indicating that mitochondrial localization of GPT activity was not necessary to meet the cell-essential demand for de novo alanine synthesis (Figure S6A).

Notably, in mediating the conversion of alanine to pyruvate, the reverse GPT reaction has been long recognized for a critical role in hepatic gluconeogenesis (Felig, 1973). RNA sequencing data from more than 50 human tissues (Genotype-Tissue Expression Project) indicate that *GPT1* is indeed most abundantly expressed in liver and has a highly restricted tissue distribution profile, but that *GPT2* expression levels are comparable among liver and several other tissues (Figure S6B). Therefore, to ask whether cellular GPT1 might instead be poised toward pyruvate formation in alanine-replete conditions, we performed analogous [U-^13^C]-alanine tracing in *GPT2*-knockout cells transduced with *GPT1* cDNA. Relative to labeling patterns measured in our control cells, the fraction of M+3-pyruvate was increased by just 1%, but that of M+3-alanine was instead markedly reduced by nearly 40%, indicating that a larger fraction of the cellular alanine pool had come from de novo synthesis as catalyzed by supraphysiologic GPT1 (Figures S6C and S6D). Remarkably, these results suggest that regardless of alanine availability, each GPT isoform was poised toward the production of alanine and αKG in this context.

Since the two human GPTs share just 67% sequence homology, we then considered if the kinetic parameters for GPT-catalyzed formation of alanine and αKG differed between the two isoforms. However, there is little biochemical characterization reported for the human GPTs, and standard GPT assays are based on indirect readouts that rely on coupled activities (Glinghammar et al., 2009; Gubern et al., 1990; McAllister et al., 2013; Ouyang et al., 2016). To address this, we developed a new GPT activity assay in which reactions containing recombinant GPT, a pyridoxal phosphate cofactor, and the substrates pyruvate and glutamate, were evaluated by using liquid chromatography-mass spectrometry (LC-MS)-based detection of αKG (Figures 4G and S6E).

Estimated *K*_M_ values for glutamate were similar between the two GPTs and equivalent to a few-fold less than the measured levels of glutamate (∼6 mM) in our K562 control cells (Figure 4H). However, the estimated *K*_M_ displayed by GPT1 for pyruvate was more than 40-fold greater than that displayed by GPT2, and comparable to the measured pyruvate concentration (∼50 μM) in the same control cells (Figure 4I). These results suggest that only GPT2 would be saturated with pyruvate in this physiologic context. Since a recent study reported only minor differences between cytosolic and mitochondrial pyruvate levels (Arce-Molina et al., 2020), the subcellular localization of human GPTs would have little impact on this rationale. Of note, given the protein banding patterns we observed for each recombinant GPT, our *k*_cat_ values for GPT2 could be underestimated compared to those for GPT1, but with no influence otherwise on measured *K*_M_ values (Figure S6F). Lastly, we expect that GPT1 and/or GPT2 can be poised to catalyze the reverse GPT reaction toward pyruvate and glutamate in either liver or other physiologic contexts that perhaps require support of distinct metabolic demands.

### Identification of a gene-nutrient interaction between *GLS* and pyruvate

We could also use our data to identify genes that encode current therapeutic targets of interest in the cancer metabolism field. For example, the glutaminase gene *(GLS)* was among the top 20 most strongly scoring RPMI-essential hits from our genome-wide screens in K562 cells (Table S1). This conditional growth phenotype was recapitulated in secondary screens but shared in only one other cell line (MOLM13), illustrating that cell-specific characteristics also contribute to context-dependent *GLS* essentiality (Figures 5A and S7A). GLS catalyzes the hydrolysis of glutamine to glutamate, an amino acid that has a number of possible downstream cellular fates, including its reversible conversion to αKG (Figure 5B) (Altman et al., 2016).

**Figure 5.**
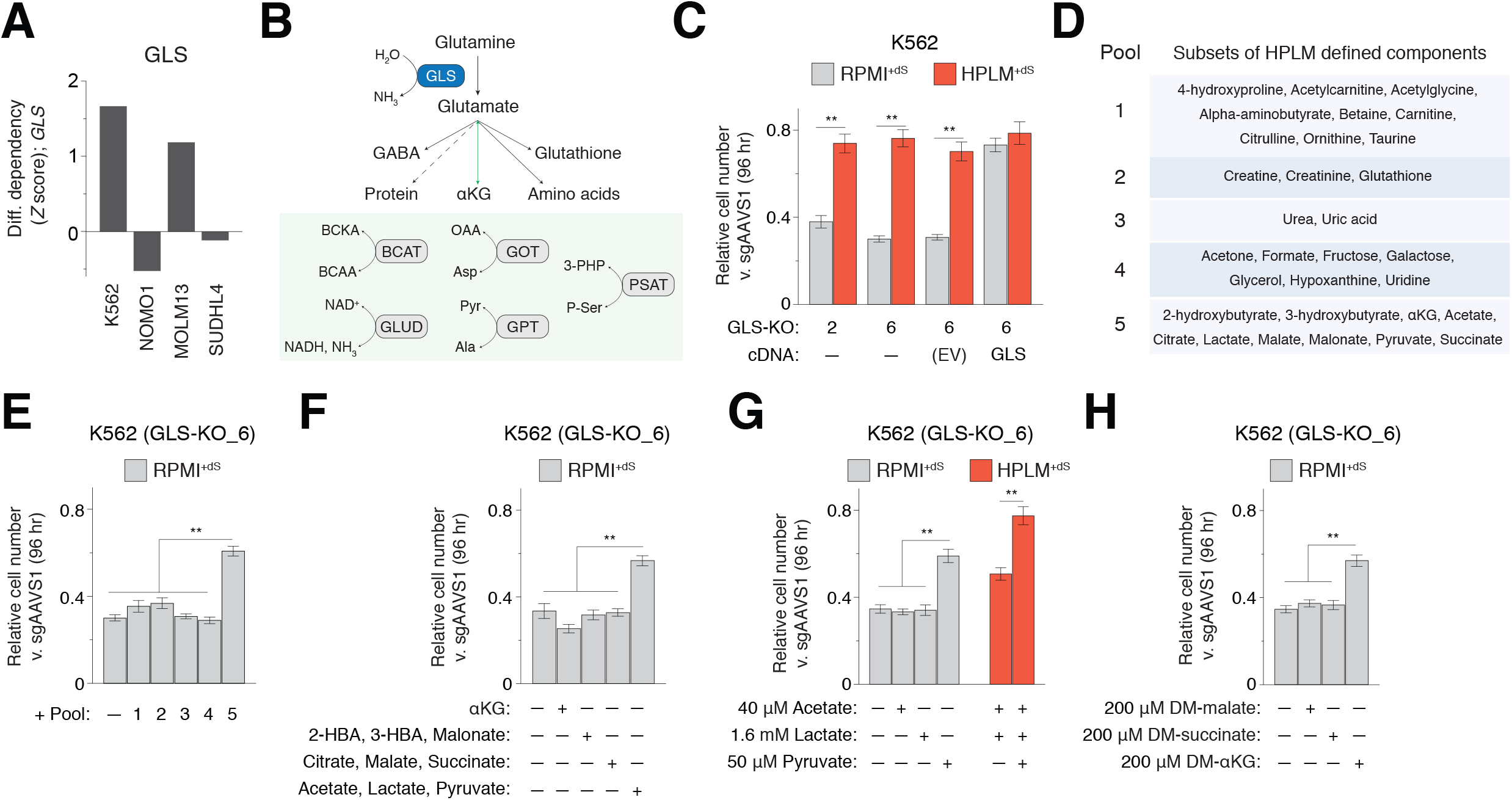
Identification of a gene-nutrient interaction between GLS and pyruvate. **See also Figure S7** (A) Conditional phenotypes for *GLS* from secondary screen results. Data for K562 are from pooled replicates. (B) Reaction catalyzed by GLS and the various cellular fates of glutamate, including its reversible conversion to αKG, green arrow (top). Several enzymes can mediate this conversion, as coupled to distinct reactions (bottom). GABA, gamma-aminobutyric acid. BCKA, branched-chain keto acid. BCAA, branched-chain amino acid. OAA, oxaloacetate. 3-PHP, 3-phosphohydroxypyruvate. P-Ser, phosphoserine. (C) Relative growth of *GLS*-knockout versus control cells in the indicated conditions (mean ± SD, *n* = 3, \*\**P* < 0.005). Knockout cells transduced with an expression construct either containing a sgRNA-resistant *GLS* (GLS) or lacking a cDNA (EV, empty vector). (D) Metabolites comprising various subsets of defined HPLM components. (E-H) Relative growth of *GLS*-knockout versus control cells in the indicated conditions (mean ± SD, *n* = 3, \*\**P* < 0.005). Pool designations correspond to those in panel D (E). Metabolites added at their HPLM-defined concentrations (F). DM-αKG, dimethyl αKG. DM-malate, dimethyl malate. DM-succinate, dimethyl succinate.

Since glutamine is an important substrate for cell growth, GLS has garnered interest as a target for cancer treatment, and recent studies report that environmental factors can influence the cellular response to GLS inhibition (Davidson et al., 2016; Muir et al., 2017). Notably, others have traced this effect to the availability of cystine (Muir et al., 2017). However, given that cystine levels between HPLM^+dS^ and RPMI^+dS^ differ by less than 2-fold, we reasoned that an alternative gene-nutrient interaction could explain the medium-dependent phenotype for loss of *GLS* suggested by our screen results (Figure S7B). To pursue this, we generated *GLS*-knockout K562 clonal cells, which as anticipated, showed a greater growth defect in RPMI^+dS^ than in HPLM^+dS^ (Figure 5C). Importantly, the expression of a sgRNA-resistant *GLS* cDNA normalized this defect between the two media, though did not fully restore relative growth compared to control cells – likely owing to clonal characteristics unrelated to GLS (Figure S7C).

Because glutamate levels in HPLM and RPMI are comparable, we then considered a more systematic testing of RPMI^+dS^ derivatives containing pools of HPLM-specific components, finding that only the addition of WSAs could effectively boost the relative growth of *GLS*-knockout cells (Figures 5D and 5E). Through additional rounds of subdivision, we ultimately pinpointed this effect to differential pyruvate availability, and in addition, found that culture in HPLM^+dS^ lacking pyruvate caused a comparable growth defect in these cells as well (Figures 5F and 5G). Since pyruvate is an upstream substrate in metabolic pathways that can generate αKG, we considered whether the *GLS* gene-nutrient interaction with pyruvate might be linked to αKG production. Interestingly, we found that adding supraphysiologic DM-αKG to RPMI^+dS^ could rescue the relative growth defect of *GLS*-knockout cells by the same extent as physiologic pyruvate, but that instead supplementing with other cell-permeable products of glutamine catabolism (DM-malate and DM-succinate) at an equivalent concentration could not (Figure 5H). Notably, a recent study described that pyruvate uptake could induce αKG formation in the context of breast cancer cells as well (Elia et al., 2019).

Our results suggest that αKG production underlies a cell-specific gene-nutrient interaction between *GLS* and pyruvate. Given that the expression of our *GLS* cDNA could boost the relative growth of *GLS-*knockout cells by a modestly greater extent (10%) than supplementation with either pyruvate or DM-αKG, it is possible that additional fates of glutamate metabolism contribute to the observed conditional *GLS* essentiality as well. Of note, the addition of physiologic pyruvate to RPMI^+dS^ had no effect on the growth defect of *GLS*-knockout cells following 24 hr culture (Figure S7D), and furthermore, the question of how pyruvate metabolism complements αKG synthesis downstream of GLS in the K562 cell line remains.

### Basal and serum components of complete culture media influence gene essentiality

Finally, given the overall prevalence of specific basal media and FBS supplements used in most CRISPR screens from DepMap, we sought to further investigate how the composition of complete media affects gene essentiality. By using our focused sgRNA library, we screened K562 cells in each of six conditions: (1) RPMI^+dS^; (2) RPMI^+S^; (3) DMEM^+dS^, 5 mM glucose; (4) DMEM^+S^, 5 mM glucose; (5) HPLM^+dS^; and (6) minimal HPLM, a modified basal HPLM that only contains glucose, salts, vitamins, and amino acids (mHPLM^+dS^) (Figure 6A, Tables S3 and S5). Of note, the screens performed in RPMI^+dS^ and HPLM^+dS^ within this scheme served as the replicates to establish our pooled datasets, and in addition, secondary screen results showed the highest correlation with those carried out in the same basal medium (Figure S8A).

**Figure 6.**
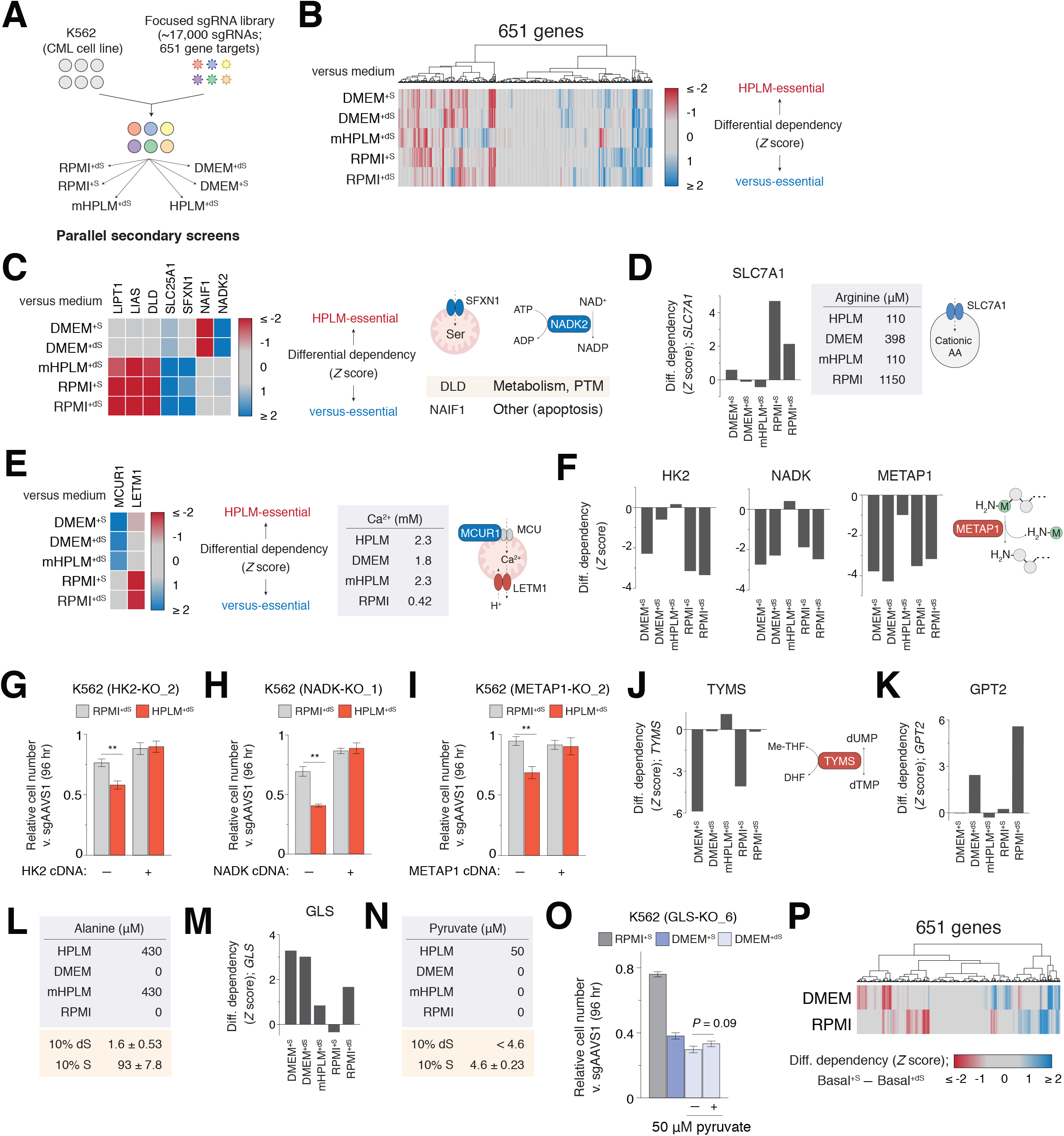
Basal and serum components of complete culture media affect gene essentiality. **See also Figures S8 and S9; Table S3** (A) Schematic for secondary screens of K562 cells in each of six media. RPMI^+S^: RPMI 1640 with 5 mM glucose and 10% FBS. DMEM^+dS^: DMEM with 5 mM glucose and 10% dialyzed FBS. DMEM^+S^: DMEM with 5 mM glucose and 10% FBS. mHPLM^+dS^: minimal HPLM and 10% dialyzed FBS (Cantor et al., 2017). (B) Cluster map showing conditional phenotypes relative to HPLM^+dS^ for genes targeted by the focused sgRNA library across six different media. Secondary screen results for each of RPMI^+dS^ and HPLM^+dS^ are from pooled replicates in all panels. Conditional phenotypes were determined as HPLM^+dS^ – “versus medium”. (C) Heatmap of conditional phenotypes for the indicated genes relative to HPLM^+dS^ (left). SFXN1 is a mitochondrial serine transporter, reaction catalyzed by NADK2, manually curated cellular processes for each of DLD and NAIF1 (right). Remaining genes are highlighted in Figure 2. (D) Conditional phenotypes for *SLC7A1* relative to HPLM^+dS^ (left). Defined arginine levels in each basal medium (middle). SLC7A1 is a plasma membrane transporter of arginine and other cationic amino acids (right). (E) Heatmap of conditional phenotypes for *MCUR1* and *LETM1* relative to HPLM^+dS^ (left). Defined Ca^2+^ availability in each basal medium (middle). LETM1 is a mitochondrial H^+^/Ca^2+^ antiporter and MCUR1 is a regulator of the MCU, mitochondrial Ca^2+^ uniporter (right). (F) Conditional phenotypes for *HK2* (left), *NADK* (middle), and *METAP1* (right) relative to HPLM^+dS^. METAP1 catalyzes the removal of N-terminal methionine from proteins (far right). (G-I) Relative growth of *HK2*-knockout (G), *NADK*-knockout (H), or *METAP1*-knockout versus control K562 cells in the indicated conditions (mean ± SD, *n* = 3, \*\**P* < 0.005). (J) Conditional phenotypes for *TYMS* relative to HPLM^+dS^ (left). Reaction catalyzed by TYMS (right). Me-THF, 5,10-methylene-5,6,7,8-tetrahydrofolate. DHF, 7,8-dihydrofolate. (K) Conditional phenotypes for *GPT2* relative to HPLM^+dS^. (L) Defined alanine concentrations in each basal medium (top). Measured alanine levels in either 10% dialyzed (dS) or untreated FBS (S) as determined by metabolite profiling of RPMI^+dS^ and RPMI^+S^, respectively (mean ± SD, *n* = 3). (M) Conditional phenotypes for *GLS* relative to HPLM^+dS^. (N) Defined pyruvate concentrations in each basal medium (top). Measured pyruvate levels in either 10% dS or S as determined by metabolite profiling of RPMI^+dS^ and RPMI^+S^, respectively (mean ± SD, *n* = 3). Pyruvate could not be readily detected in RPMI^+dS^ by the applied method. (O) Relative growth of *GLS*-knockout versus control cells in the indicated conditions (mean ± SD, *n* = 3, \*\**P* < 0.005). (P) Cluster map showing differential dependency calculated as DMEM^+S^ – DMEM^+dS^ (top row) or RPMI^+S^ – RPMI^+dS^ (bottom row).

Next, we standardized the differential gene scores between HPLM^+dS^ and each of the five other conditions, and then found that the sets of genes differentially essential in either HPLM (negative) or the relative medium (positive) showed varying degrees of overlap and a number of distinct patterns (Figure 6B). For example, we identified *ACLY* as a positive hit gene in all five relative conditions, again likely owing to the defined availability of acetate in HPLM (Figure S8B). In contrast, the putative diphosphate kinase *NME6* instead scored as HPLM-essential in each comparison, but the cause for this conditional growth phenotype is unknown (Figure S8C).

Conditional phenotypes of other genes were instead common to either non-DMEM *(DLD* and *SFXN1)* or DMEM-based media *(NADK2* and *NAIF1*) (Figure 6C). *SLC7A1*, a cationic amino acid transporter gene, scored as a positive hit only in RPMI-based media, perhaps indicative of an increased dependence on arginine uptake driven by culture in highly supraphysiologic arginine conditions (Figure 6D). Similarly, HPLM-relative phenotypes of genes involved in Ca^2+^ transport *(LETM1* and *MCUR1)* might be explained by the differential Ca^2+^ availability in RPMI versus both HPLM and DMEM (Figure 6E).

Our analysis also revealed hits common to all conditions except for mHPLM^+dS^, suggesting conditional phenotypes attributed to concentration differences among the amino acids and salt ions in HPLM compared to each of RPMI and DMEM (Figure S8D). Among these hit genes were *HK2, NADK*, and a methionine aminopeptidase *(METAP1)* (Figure 6F). Notably, we generated knockout clonal cells and sgRNA-resistant cDNAs to validate the HPLM-essential phenotypes for each of these by using relative growth assays in HPLM^+dS^ versus RPMI^+dS^, though the associated gene-nutrient interactions are unknown (Figures 6G, 6H, 6I, S8E, S8F, and S8G). We also found that *TBC1D31*, a gene of unknown function, was HPLM-essential only relative to mHPLM^+dS^, suggesting a medium-dependent phenotype that further requires the physiologic availability of additional components among the amino acids and salt ions in HPLM (Figure S8H).

Interestingly, we identified thymidylate synthase *(TYMS)* as a negative hit compared only to each of RPMI^+S^ and DMEM^+S^, suggesting that 10% untreated FBS provides an otherwise undefined component that may support pyrimidine metabolism (Figure 6J). Moreover, while genes associated with the UFMylation machinery scored as either positive hits in RPMI^+dS^ or negative hits in DMEM^+dS^, their conditional phenotypes were largely diminished in comparisons made to complete media counterparts prepared with untreated FBS (Figure S8I).

We next investigated if conditional phenotype patterns for *GPT2* and *GLS* were consistent with gene-nutrient interactions identified above. First, we found that *GPT2* was a strongly positive hit only in RPMI^+dS^ and DMEM^+dS^ (Figures 6K and S9A). Notably, while alanine is not defined in either RPMI or DMEM, 10% untreated versus dialyzed FBS contributes alanine at a more than 50-fold greater concentration equivalent to just less than 4-fold that defined in HPLM (Figure 6L). Consistent with our screen results, *GPT2*-knockout cells did not exhibit a growth defect in either RPMI^+S^ or DMEM^+S^, indicating that the sub-physiologic alanine in these two media was sufficient to complement *GPT2* deletion (Figure S9B). Further, as expected, these cells also showed a growth impairment in DMEM^+dS^ that could be fully rescued by the addition of physiologic alanine. Of note, conditional phenotype patterns for each of *MPC1, MPC2*, and *AARS* largely reflected that of *GPT2* as well (Figures S9C, S9D, and S9E). Second, we identified *GLS* as a positive hit in RPMI^+dS^, mHPLM^+dS^, and each DMEM-based medium (Figures 6M and S9F). Surprisingly, however, *GLS* did not show a strong conditional phenotype versus RPMI^+S^ which, like DMEM^+S^, contains pyruvate from untreated FBS at a concentration 10-fold lower than in HPLM (Figure 6N). Indeed, we then observed that *GLS* deletion caused growth defects that were comparable either between RPMI^+S^ and HPLM^+dS^ or instead between the two DMEM-based media and RPMI^+dS^, and further, that adding 50 μM pyruvate to DMEM^+dS^ had little impact on the relative growth of these cells (Figure 6O). Remarkably, these results indicate that the *GLS* gene-nutrient interaction with pyruvate in K562 cells is itself context-dependent, as physiologic pyruvate was neither necessary nor sufficient to largely complement *GLS*-knockout in all media conditions. Furthermore, cystine levels in DMEM are equivalent to those in RPMI (Figure S9G).

Our analysis also revealed other genes with HPLM-relative phenotypes that were similarly unique to a specific RPMI- or DMEM-based medium, including hits common among all conditions except for either DMEM^+S^ *(MTHFD1L, MTHFD2)* or RPMI^+S^ *(KEAP1)*, and another that showed a positive growth phenotype relative only to RPMI^+S^ *(TAF10)* (Figure S8J). Indeed, the overall sets of differential dependencies calculated between either the two RPMI-based or the two DMEM-based media showed little overlap (Figure 6P), illustrating that the undefined contents of untreated FBS can have strikingly disparate effects on gene essentiality depending on the basal medium supplemented.

## CONCLUSIONS

Conditional gene essentiality profiling in human blood cancer cell lines revealed hundreds of medium-dependent fitness genes that span several cellular processes and vary with natural cell-intrinsic diversity, suggesting that forward genetic screens in HPLM should make it possible to define new cancer-specific vulnerabilities and gene-nutrient interactions in human cells.

CRISPR screens in standard media conditions that consist of a basal medium with little relevance to human blood and an untreated FBS supplement have likely masked many genetic dependencies in human cancers. Here we identify strong loss-of-function phenotypes for *GPT2* and genes encoding the MPC. We traced these effects to the differential availability of alanine – one of three GPT reaction components uniquely defined in HPLM versus both RPMI and DMEM. By mediating de novo alanine production from mitochondrial pyruvate imported via the MPC, we find that GPT2 supports the cell-essential demand of protein synthesis in conditions of relative alanine limitation. Notably, traditional basal media such as RPMI and DMEM are most often supplemented with 10% untreated FBS which, among countless undefined components, provides alanine at sub-physiologic levels that are nonetheless sufficient to complement *GPT2* deletion.

Here we have also uncovered a cell-specific conditional phenotype for loss of *GLS*, which encodes a current target of interest for cancer therapy. However, glutamate levels are comparable between HPLM and RPMI, and though others have shown that cystine availability can influence cell response to GLS inhibition, cystine levels in HPLM versus each of RPMI and DMEM differ by less than 2-fold. Therefore, by applying a systematic approach, we traced the loss-of-function phenotype for *GLS* to a gene-nutrient interaction with pyruvate, another metabolite undefined in both RPMI and DMEM. We find that this interaction is associated with the generation of αKG, one of several fates of cellular glutamate. Remarkably, however, we found that conditional phenotypes for *GLS* deletion in K562 cells could not be explained by relative pyruvate availability in all media conditions, offering added complexity into understanding context-dependent *GLS* essentiality in human cancers. We also identified other genes with strong HPLM-relative phenotypes specific to a particular RPMI- or DMEM-based medium, further suggesting that reported genetic dependency data across hundreds of human cancer lines has been influenced in part by the conventional media conditions used for individual forward genetic screens.

Efforts to uncover the gene-nutrient interactions for most hits from our screens will require unbiased and systematic approaches as used for *GLS*. For example, we also validated conditional phenotypes for each of three HPLM-essential hit genes *(HK2, NADK*, and *METAP1)*, though the underlying gene-nutrient interactions are not yet known. However, our screen results do suggest that these could be traced to medium components among the amino acids and salt ions. Of note, all three genes are co-expressed with functionally related homologs in most cancer cell lines and, interestingly, *NADK* is not annotated as an essential gene in any of the nearly 800 reported CRISPR screens in DepMap.

Medium composition is a relatively flexible and accessible environmental factor among those that can influence gene essentiality. These attributes should make it possible to investigate how the genetic drivers of selectable phenotypes in human cells (e.g. growth, cell state and drug response) might vary with different biochemical conditions, including potential HPLM derivatives or physiologic media designed to better recapitulate other biofluids (Cantor, 2019). By performing comparative CRISPR screens based on medium composition, it may also be possible to design cancer treatment strategies that exploit gene-nutrient interactions by coupling targeted inhibitors with either dietary or enzyme-catalyzed modulation of circulating metabolites. Notably, a number of clinical and preclinical enzymes mediate systemic depletion of specific metabolites (Cantor and Sabatini, 2012; Cantor et al., 2012; Cramer et al., 2016; Lu et al., 2020; Patgiri et al., 2020; Triplett et al., 2018), and recent studies have shown that dietary intervention to either restrict or increase the availability of specific amino acids can affect cancer growth and chemotherapeutic efficacy (Gao et al., 2019; Kanarek et al., 2018).

## SUPPLEMENTAL INFORMATION

Supplemental information includes all methods, nine figures, and six tables.

## ACKNOWLEDGEMENTS

We thank members of the Sabatini lab for helpful insights and members of the Cantor lab for maintaining the LC-MS platform. We also thank S. Gupta and P. Thiru for assistance with the preparation and analysis of RNA sequencing data, as well as O. Wurtzel for initial sequencing verification of the focused sgRNA library. This work was supported by grants from the NIH (R01CA103866 and R37AI047389 to D.M.S. and K22CA225864 to J.R.C.) and the Koch Institute (Frontier Research Grant) to D.M.S. N.J.R. and J.R.C. were also supported by startup funds from the Morgridge Institute for Research. Fellowship support was provided by the NIH (T32HG002760 to K.S.H., F31CA228241-01 to C.H.A., and T32GM008349 to R.W.S.). D.M.S. is an investigator of the Howard Hughes Medical Institute and an American Cancer Society Research Professor.

## AUTHOR CONTRIBUTIONS

J.R.C. and D.M.S. initiated the project. J.R.C. designed the research plan, conducted the screens, performed the experiments with assistance from K.S.H and R.W.S., and analyzed experimental data. N.J.R. analyzed the screen data with assistance from C.H.A., H.R.K. and J.R.C. C.H.A. designed the focused sgRNA library. J.R.C. wrote the manuscript, H.R.K. and D.M.S. provided editing input.

## SUPPLEMENTAL INFORMATION

### Supplemental Figure Legends

**Figure S1.**
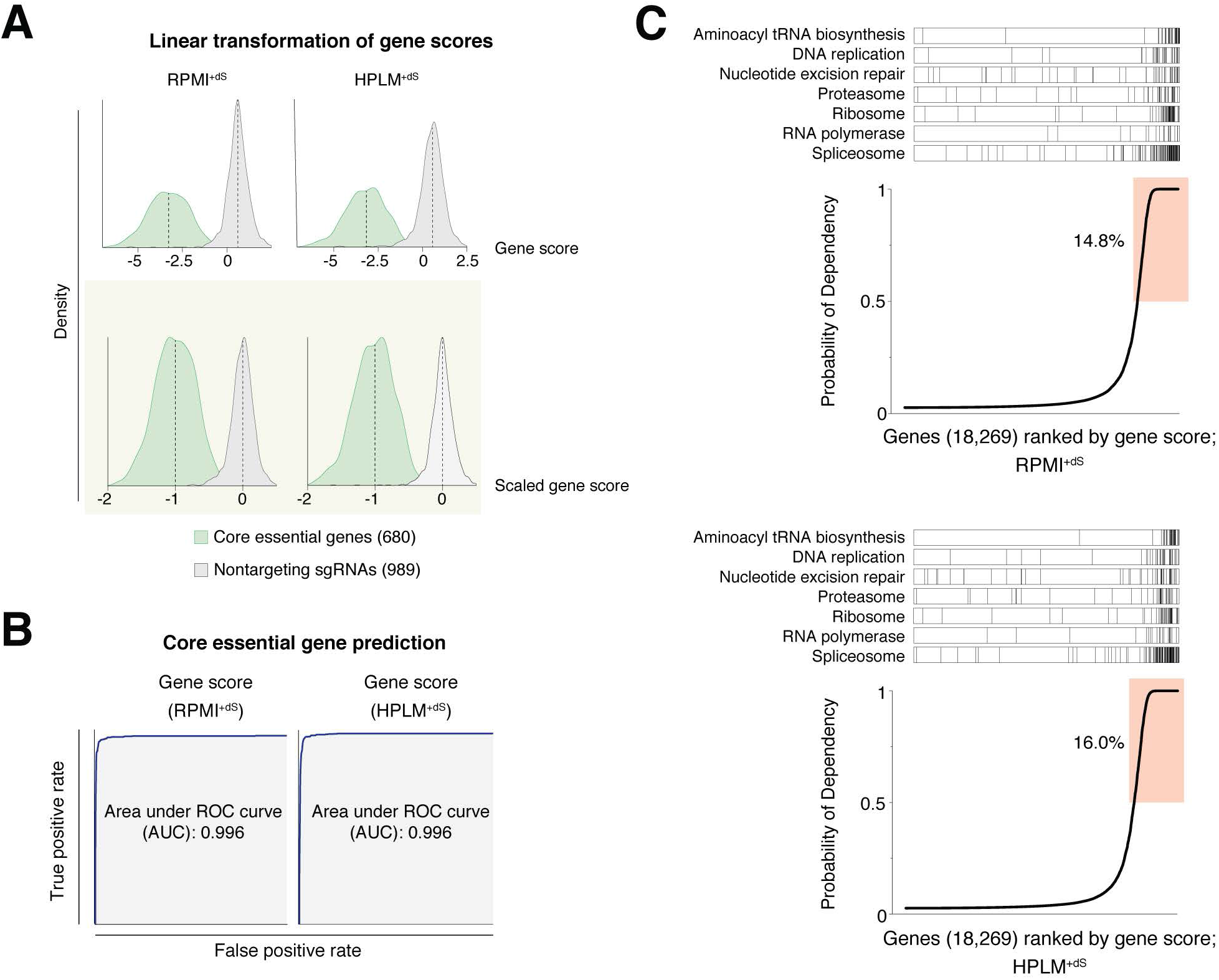
Gene score scaling and genome-wide screen performance. **Related to Figure 1** (A) Raw gene score distributions for 680 CEGs (green) and 989 non-targeting sgRNAs (gray) from genome-wide screens in RPMI^+dS^ (left) and HPLM^+dS^ (right). Distributions for each following linear transformation scaling (bottom). (B) Receiver operator characteristic (ROC) curves for the prediction of CEGs using datasets from genome-wide screens in RPMI^+dS^ (left) and HPLM^+dS^ (right). (C) Plots of all targeted genes ranked by probability of dependency from genome-wide screens in RPMI^+dS^ (top) and HPLM^+dS^ (bottom). Shaded regions indicate a probability greater than 0.5. Barcode plots depict the distribution of genes involved in fundamental cellular processes.

**Figure S2.**
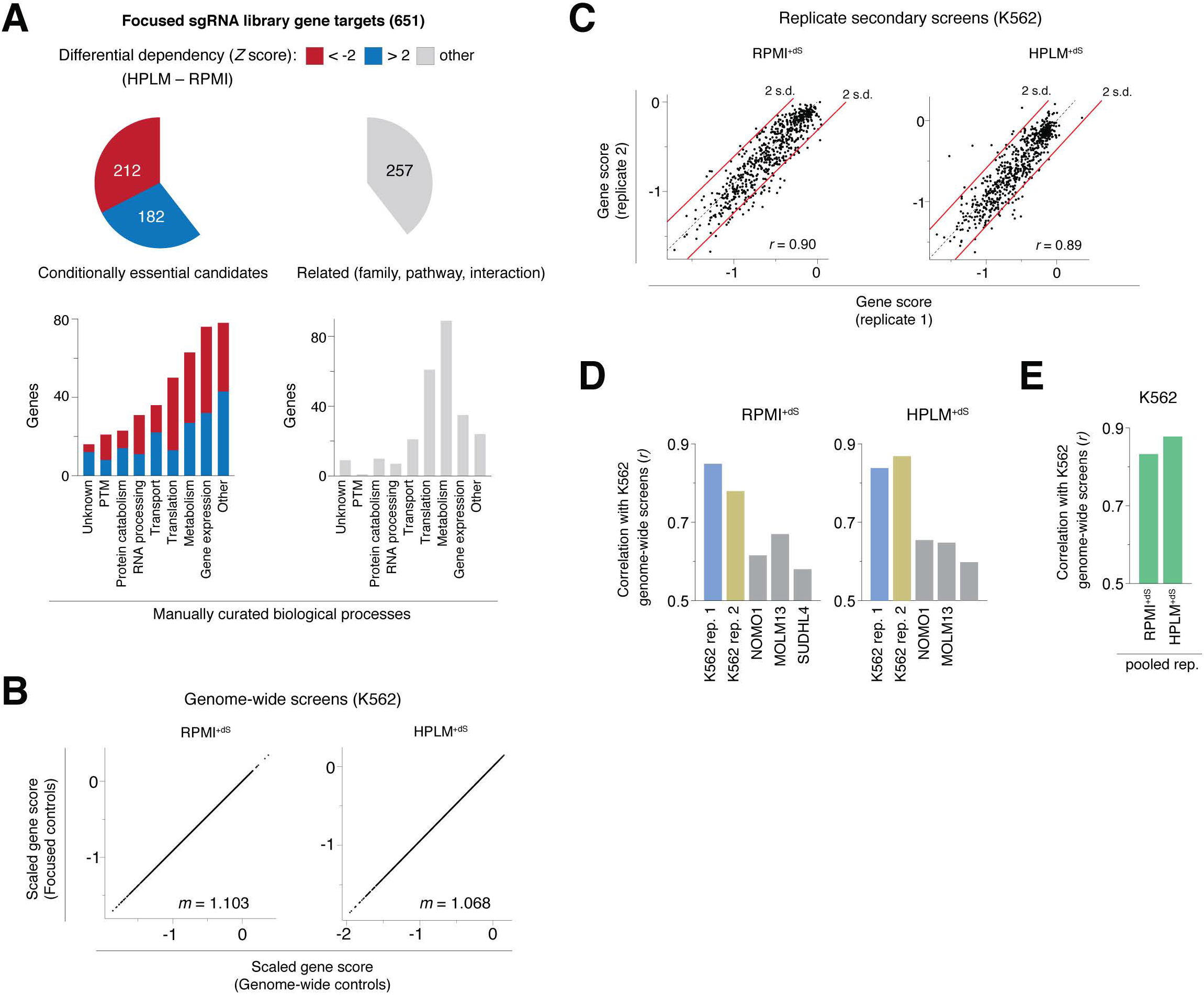
Focused sgRNA library design and correlation analyses. **Related to Figure 2** (A) Pie chart depicting the fraction of genes targeted by the focused sgRNA library with indicated conditional phenotypes from genome-wide screen results (top). Composition of focused sgRNA library gene targets by manually curated cellular processes (bottom). (B) Comparison of genome-wide screen results in RPMI^+dS^ (left) and HPLM^+dS^ (right) scaled using the CEG and non-targeting sgRNA control sets from either the genome-wide or focused sgRNA libraries. Each point represents one gene *m*, slope. (C) Comparison of gene scores from replicate secondary screens of K562 cells in RPMI^+dS^ (left) and HPLM^+dS^ (right). Each point represents one gene. Red lines indicate differences of greater than 2 s.d. *r*, Pearson’s correlation coefficient. (D) Gene score correlations between secondary screens of four cell lines and the genome-wide screens of K562 in each of RPMI^+dS^ (left) and HPLM^+dS^ (right). (E) Gene score correlations between the pooled K562 secondary screen dataset and genome-wide screens of K562 in each of RPMI^+dS^ and HPLM^+dS^.

**Figure S3.**
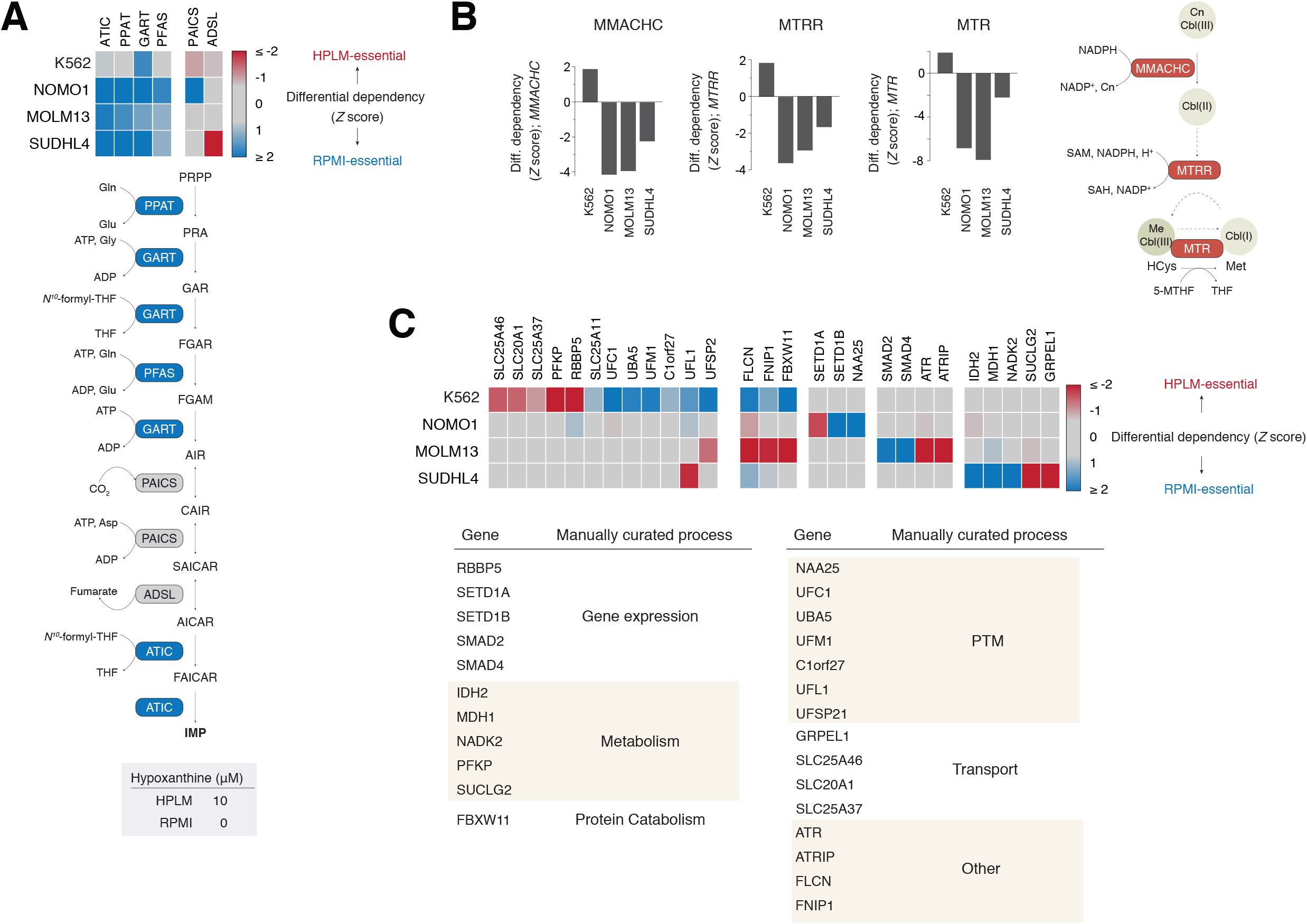
Additional examples of genes with cell-specific medium-dependent phenotypes. **Related to Figure 2** (A) Heatmap of conditional phenotypes for the indicated genes (top). Schematic of the de novo purine biosynthesis pathway (middle). Enzymes encoded by the four genes that share a common RPMI-essential phenotype in three cell lines are shaded blue and catalyze irreversible steps in the pathway based on annotations in the KEGG Pathway database. Defined hypoxanthine concentrations in HPLM and RPMI (bottom). Secondary screen data for K562 are from pooled replicates in all panels. (B) Conditional phenotypes for *MMACHC* (left), *MTRR* (middle), and *MTR* (right) from secondary screen results (left). Schematic depicting the processing of CnCbl (III) to Cbl (II) and the conversion of HCys to methionine coupled to the reductive generation of Cbl(I) to MeCbl (III) (far right). CnCbl (III), cyanocob (III)alamin; Cbl (II), cob (II)alamin; HCys, homocysteine; Cbl(I), cob(I)alamin; MeCbl (III), methylcob (III)alamin. (C) Heatmap of conditional phenotypes for the indicated genes from secondary screen results (top). Manually curated biological processes for each indicated gene (bottom).

**Figure S4.**
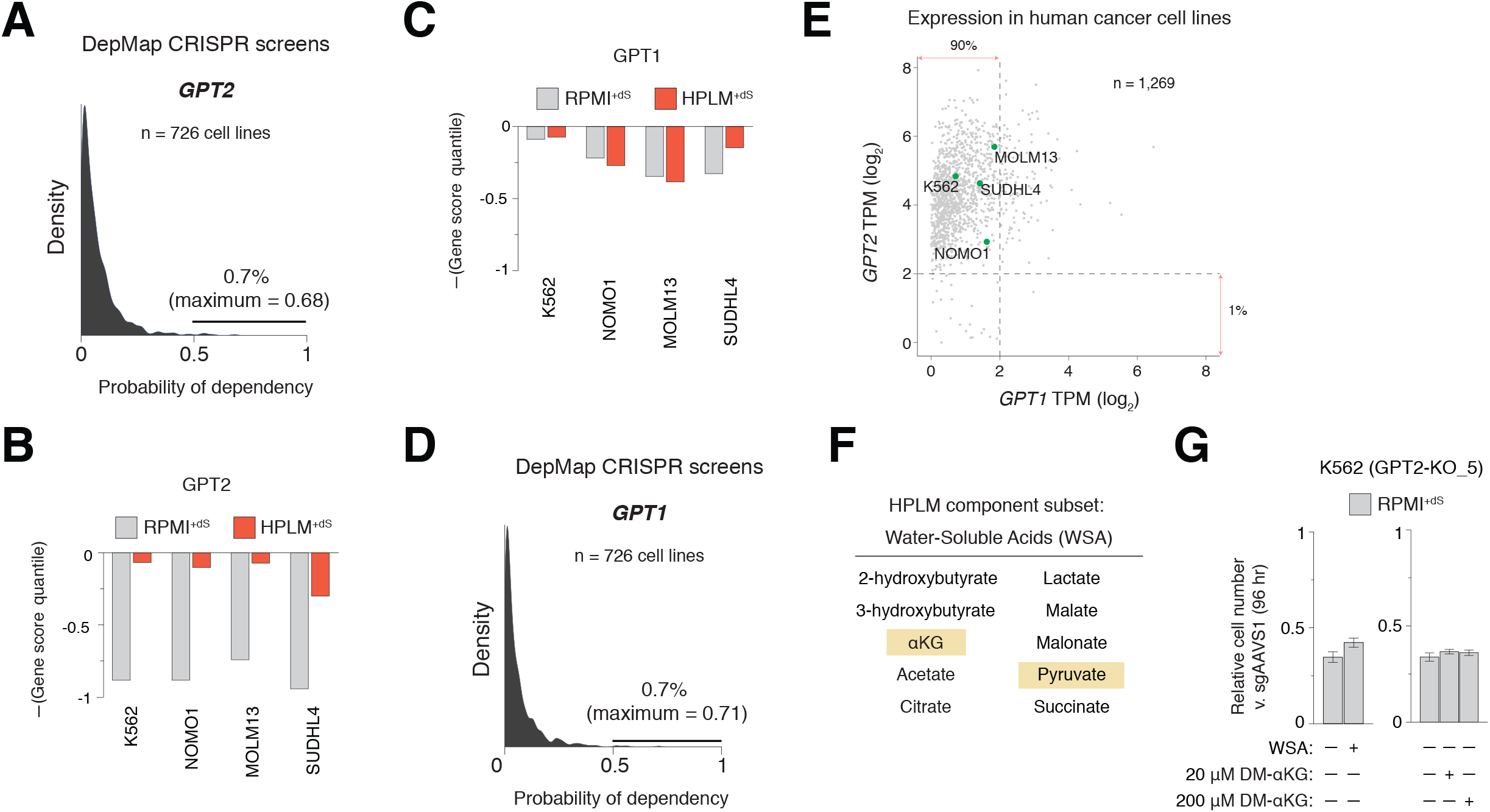
Annotated dependency and expression data for *GPT1/2* in human cancer cell lines; Quantile scores for *GPT1/2* from secondary screens; and Neither HPLM-defined water-soluble acids nor cell-permeable αKG can rescue the relative growth defect of *GPT2*-knockout cells in RPMI^+dS^. **Related to Figure 3** (A) Density plot for probability of dependency values annotated for *GPT2* from reported DepMap CRISPR screens in human cancer cell lines. Probability greater than 0.5 is the reference threshold for essentiality. (B-C) Secondary screen gene score quantiles for *GPT2* (B) and *GPT1* (C) in each medium. Data for K562 are from pooled replicates. (D) Density plot for probability of dependency values annotated for *GPT1* from reported DepMap CRISPR screens in human cancer cell lines. (E) Comparison of *GPT1* versus *GPT2* expression levels in human cancer cell lines from RNA sequencing data in DepMap. Labeled points indicate screened cell lines in this study. TPM, transcripts per million. (F) Metabolites comprising the water-soluble acids (WSAs) subset of defined HPLM components. (G) Relative growth of *GPT2*-knockout versus control cells in the indicated conditions (mean ± SD, *n* = 3, \*\**P* < 0.005). DM-αKG, dimethyl αKG.

**Figure S5.**
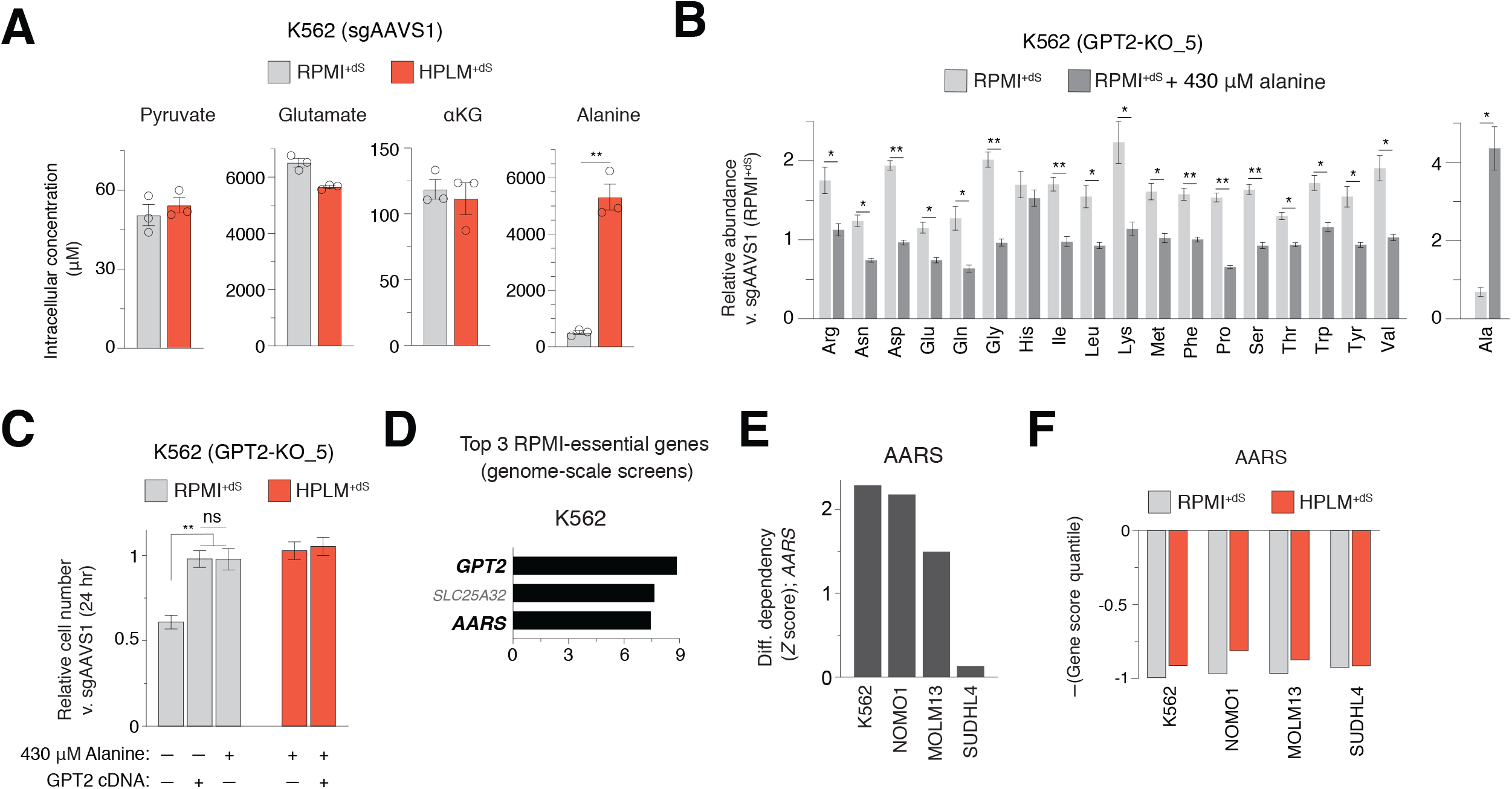
Additional data related to unbiased metabolite profiling in *GPT2*-knockout and control cells; Physiologic alanine rescues the growth defect of *GPT2*-knockout cells after 24 hr culture; and *AARS* is a strong RPMI-essential hit gene. **Related to Figure 4** (A) Intracellular concentrations of each GPT reaction component in control (sgAAVS1) K562 cells following 24 hr culture in the indicated conditions (mean ± SD, *n* = 3, \*\**P* < 0.01). (B) Relative amino acid abundances in *GPT2*-knockout cells following 24 hr culture in either RPMI^+dS^ (light gray) or RPMI^+dS^ containing 430 μM alanine (dark gray) versus those from control cells in RPMI^+dS^ (mean ± SEM, *n* = 3, \**P* < 0.05, \*\**P* < 0.005). (C) Relative growth of *GPT2*-knockout versus control cells in the indicated conditions (mean ± SD, *n* = 3, \*\**P* < 0.005). ns, not significant. (D) Top three most strongly scoring RPMI-essential hits from genome-wide screens in K562. (E) Conditional phenotypes for loss of *AARS* from secondary screen results. Data for K562 are from pooled replicates. (F) Secondary screen gene score quantiles for *AARS* in each medium.

**Figure S6.**
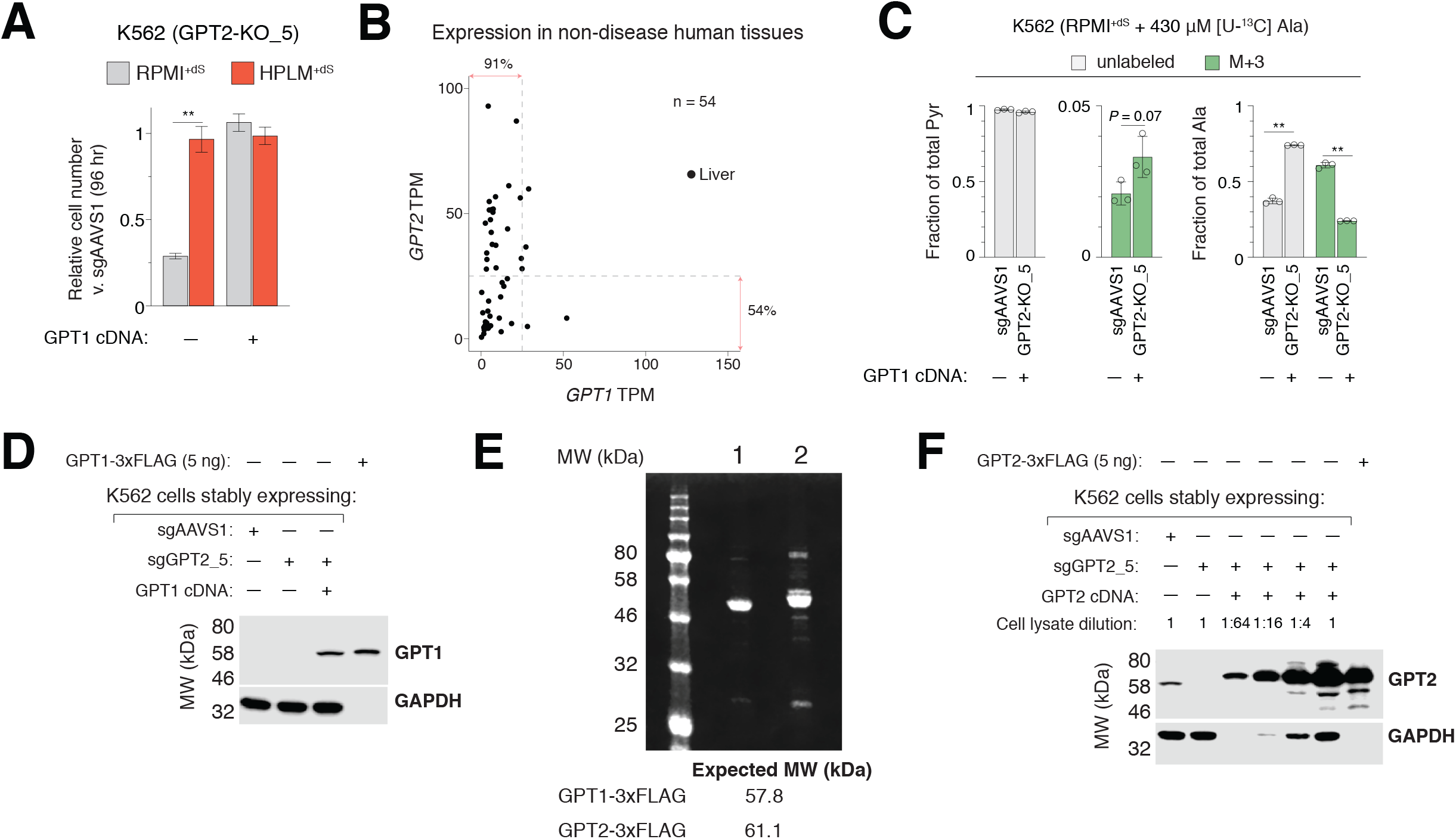
Relative growth and alanine tracing data for *GPT2*-knockout cells transduced with a *GPT1* cDNA; Annotated expression data for *GPT1/2* in normal human tissues; and Purification of recombinant GPTs. **Related to Figure 4** (A) Relative growth of *GPT2*-knockout versus control cells in the indicated conditions (mean ± SD, *n* = 3, \*\**P* < 0.005). (B) Comparison of *GPT1* versus *GPT2* expression levels in non-diseased human tissues from RNA sequencing data in GTEx. Point indicating liver is indicated. TPM, transcripts per million. (C) Fractional labeling of pyruvate (left) and alanine (right) following 24 hr culture of cells in RPMI^+dS^ containing 430 μM [U-^13^C]-alanine (mean ± SD, *n* = 3, \*\**P* < 0.005). M+3, incorporation of three ^13^C. (D) Immunoblot for expression of GPT1 in cell lysates and from sample of purified GPT1-3xFLAG. M.W. standards are annotated. GAPDH served as a loading control. (E) Pseudocolor Coomassie stained gel imaged using a LICOR Odyssey FC. 1: GPT1-3xFLAG, 2: GPT2-3xFLAG (see Methods for note on observed M.W. compared to immunoblots of the same purified proteins). (F) Immunoblot for expression of GPT2 in cell lysates and from sample of purified GPT2-3xFLAG. M.W. standards are annotated. Cells transduced with cDNA were loaded in serial dilution as indicated. GAPDH served as a loading control.

**Figure S7.**
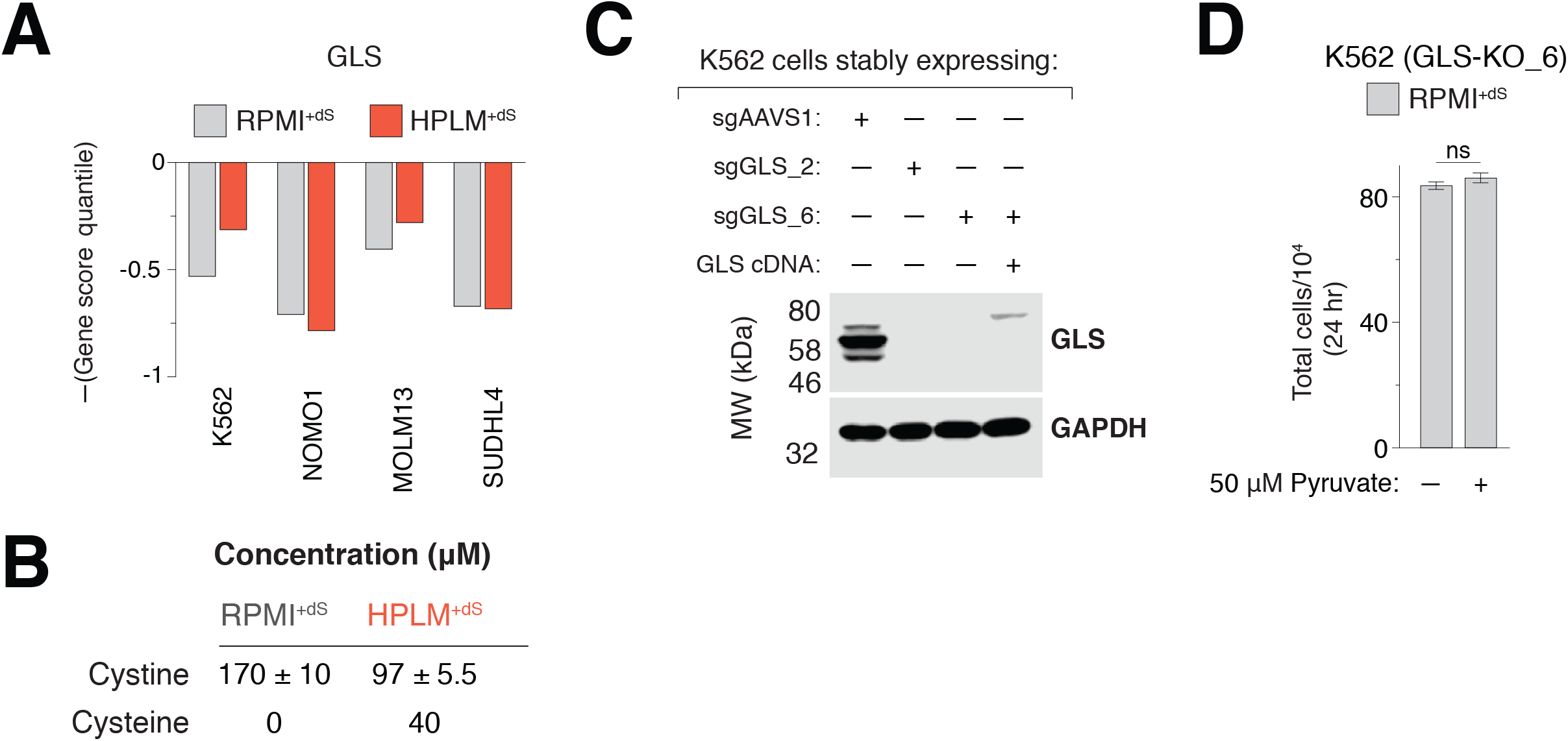
Additional data related to the conditional loss-of-function phenotype for *GLS*. **Related to Figure 5** (A) Secondary screen gene score quantiles for *GLS* in each medium. Data for K562 are from pooled replicates. (B) Cystine concentrations in RPMI^+dS^ and HPLM^+dS^ as measured by metabolite profiling (mean ± SD, *n* = 3). Cysteine cannot be readily detected in samples with the applied method but is also a defined component in HPLM. (C) Immunoblot for expression of GLS in the indicated cells. M.W. standards are annotated GAPDH served as a loading control. (D) Growth of *GLS*-knockout cells in the indicated conditions. ns, not significant.

**Figure S8.**
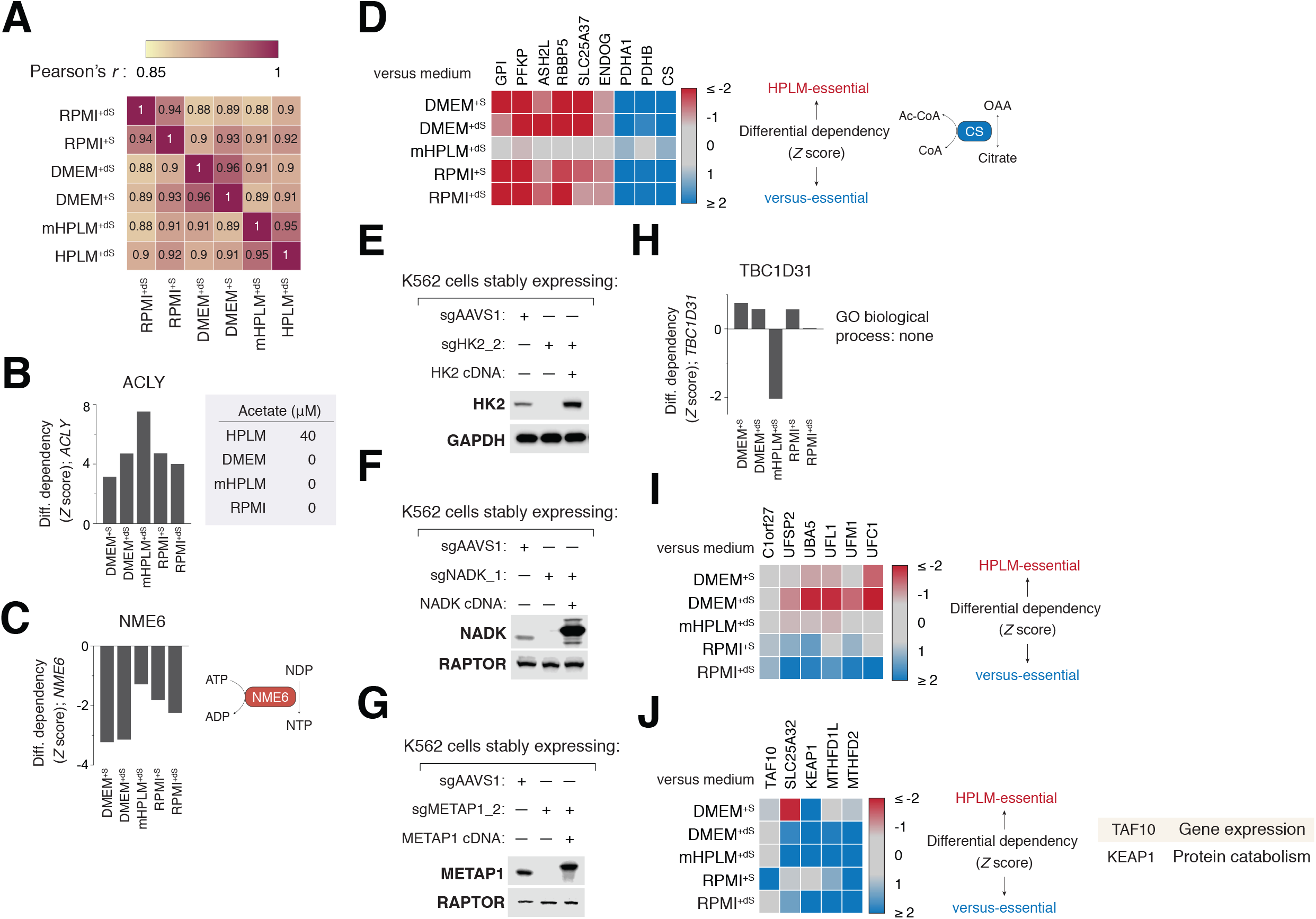
Additional examples of genes with conditional loss-of-function phenotypes and Immunoblots for expression of validated HPLM-essential hit genes. **Related to Figure 6** (A) Gene score correlations between secondary screens in K562 across six conditions. Data for RPMI^+dS^ and HPLM^+dS^ are from pooled replicates in all panels. (B-C) Conditional phenotypes for *ACLY* (B) and *NME6* (C) relative to HPLM^+dS^. Defined acetate levels in each basal medium (B, right). Reaction catalyzed by NME6 (C, right). (D) Heatmap of conditional phenotypes for the indicated genes relative to HPLM^+dS^ (left). Reaction catalyzed by CS (right). Remaining genes are highlighted elsewhere in Figure 2. (E-G) Immunoblots for expression of HK2 (E), NADK (F), and METAP1 (G) in the indicated cells. GAPDH (E) and RAPTOR (F and G) served as loading controls. (H) Conditional phenotypes for *TBC1D31*, a gene of unknown function, relative to HPLM^+dS^. (I) Heatmap of conditional phenotypes for the UFMylation machinery genes relative to HPLM^+dS^. (J) Heatmap of conditional phenotypes for the indicated genes relative to HPLM^+dS^ (left). Manually curated cellular processes for both TAF10 and KEAP1 (right). Remaining genes are highlighted in Figure 2.

**Figure S9.**
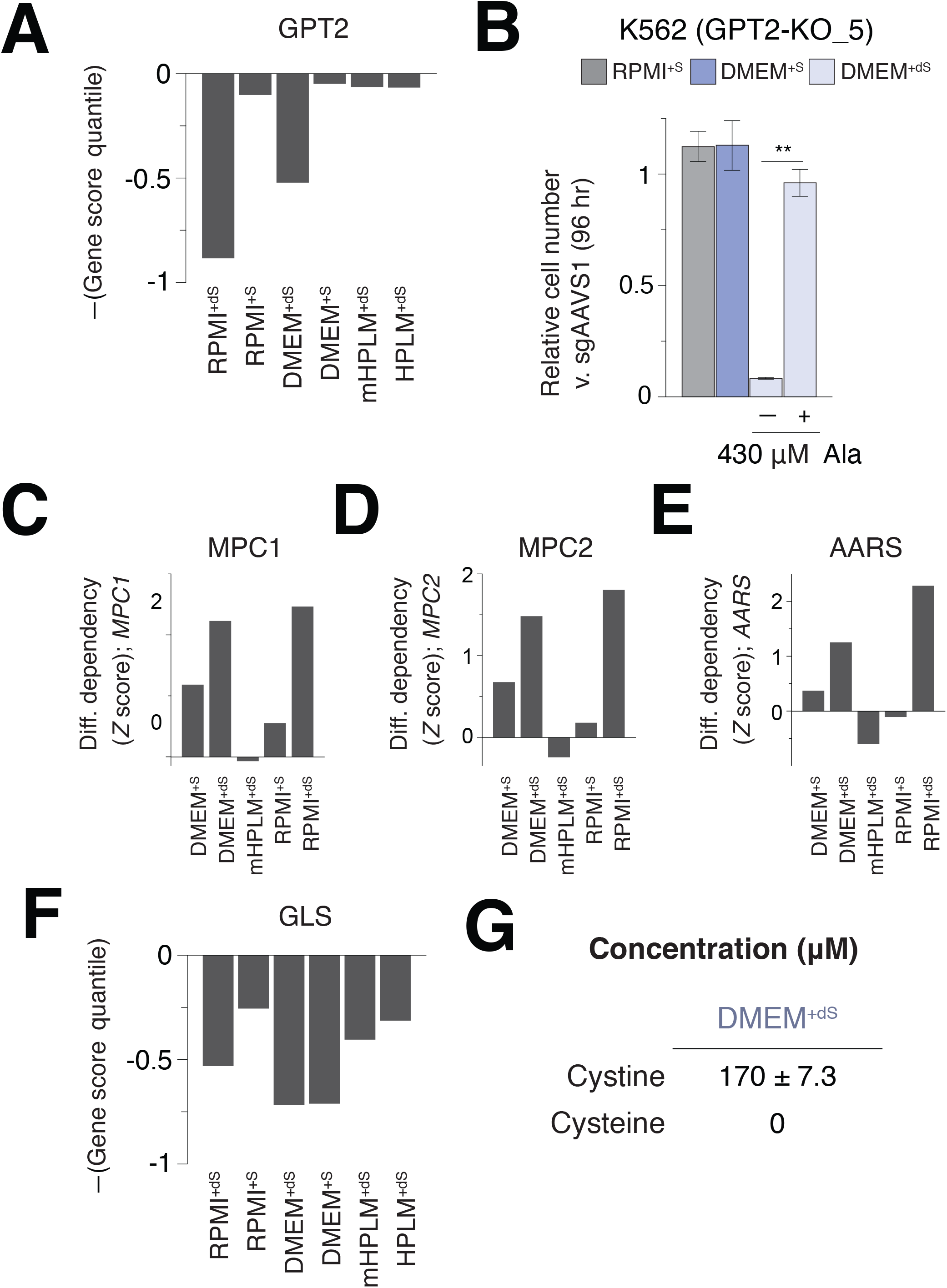
Differential dependency data for *MPC1/2* and AARS; Quantile scores for *GPT2* and *GLS* from secondary screens in K562 cells; and Sub-physiologic alanine provided by 10% untreated FBS can complement *GPT2* deletion. **Related to Figure 6** (A) Secondary screen gene score quantiles for *GPT2* in each medium. Data for RPMI^+dS^ and HPLM^+dS^ are from pooled replicates in all panels. (B) Relative growth of *GPT2*-knockout versus control cells in the indicated conditions (mean ± SD, *n* = 3, \*\**P* < 0.005). (C-E) Conditional phenotypes for *MPC1* (C), *MPC2* (D), and *AARS* (E) relative to HPLM^+dS^. (F) Secondary screen gene score quantiles for *GLS* in each medium. (G) Cystine concentration in DMEM^+dS^ as measured by metabolite profiling (mean ± SD, *n* = 3). Cysteine is not a defined component in DMEM.

### Reagents and Resources

#### Antibodies

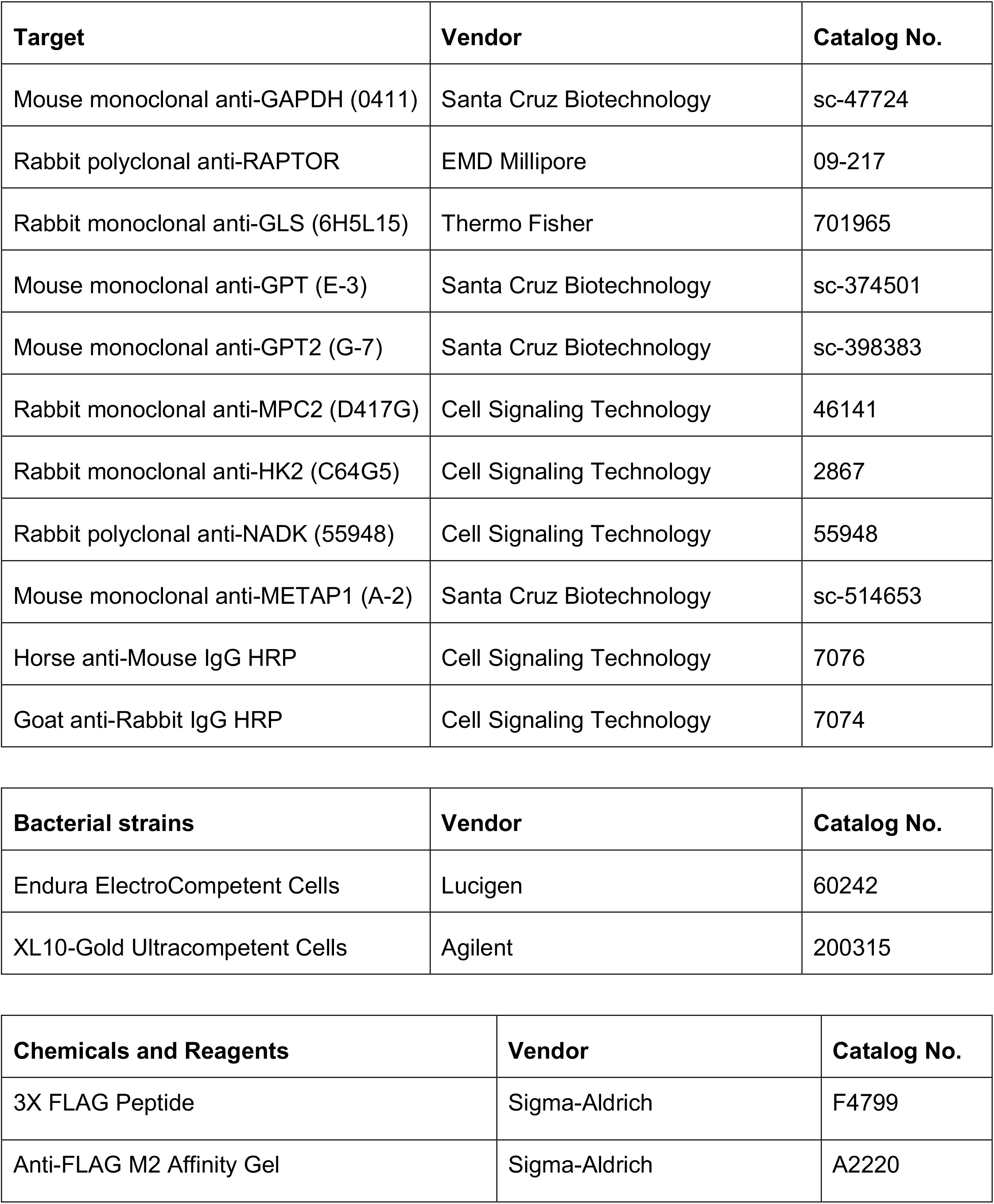
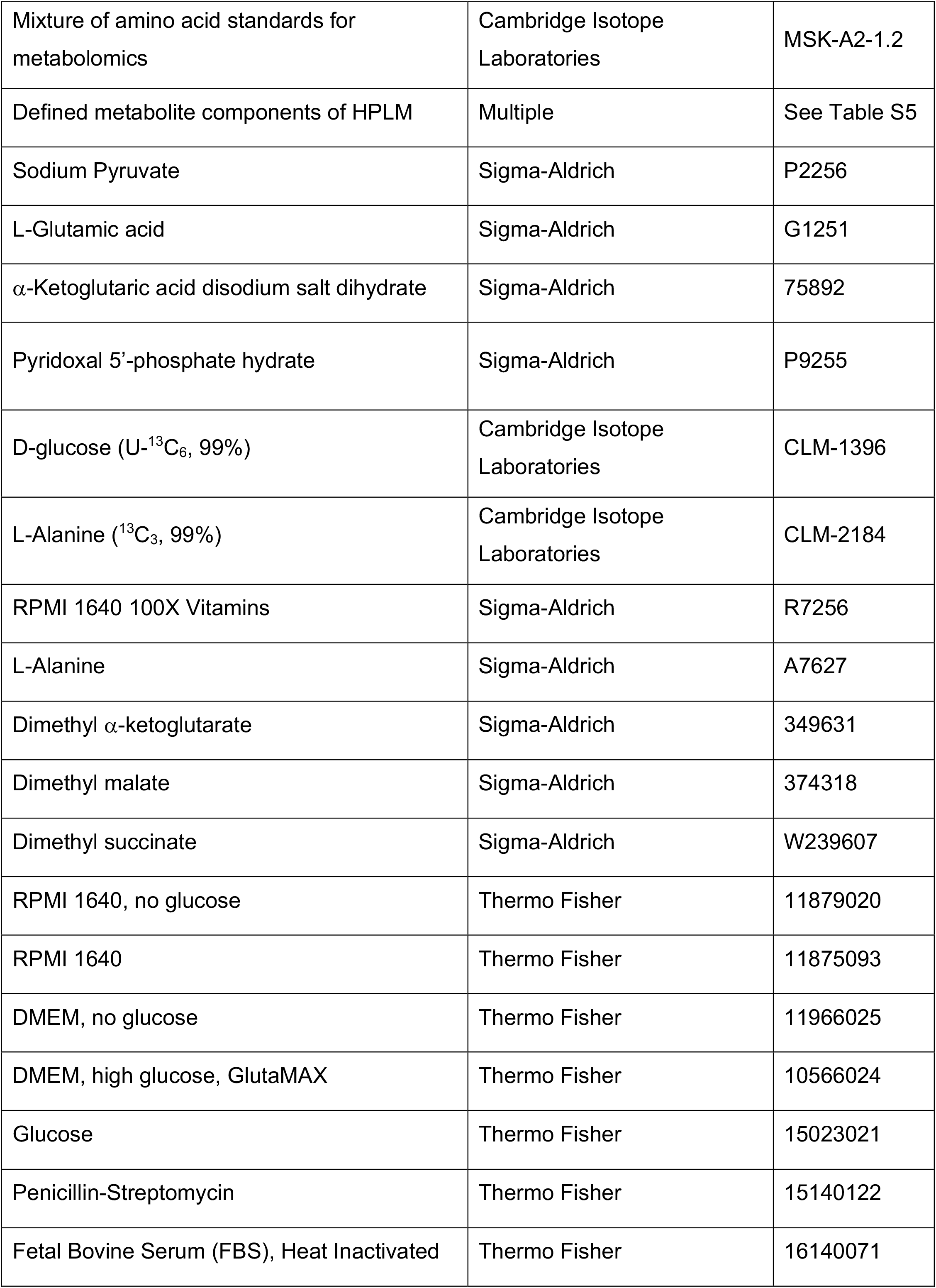
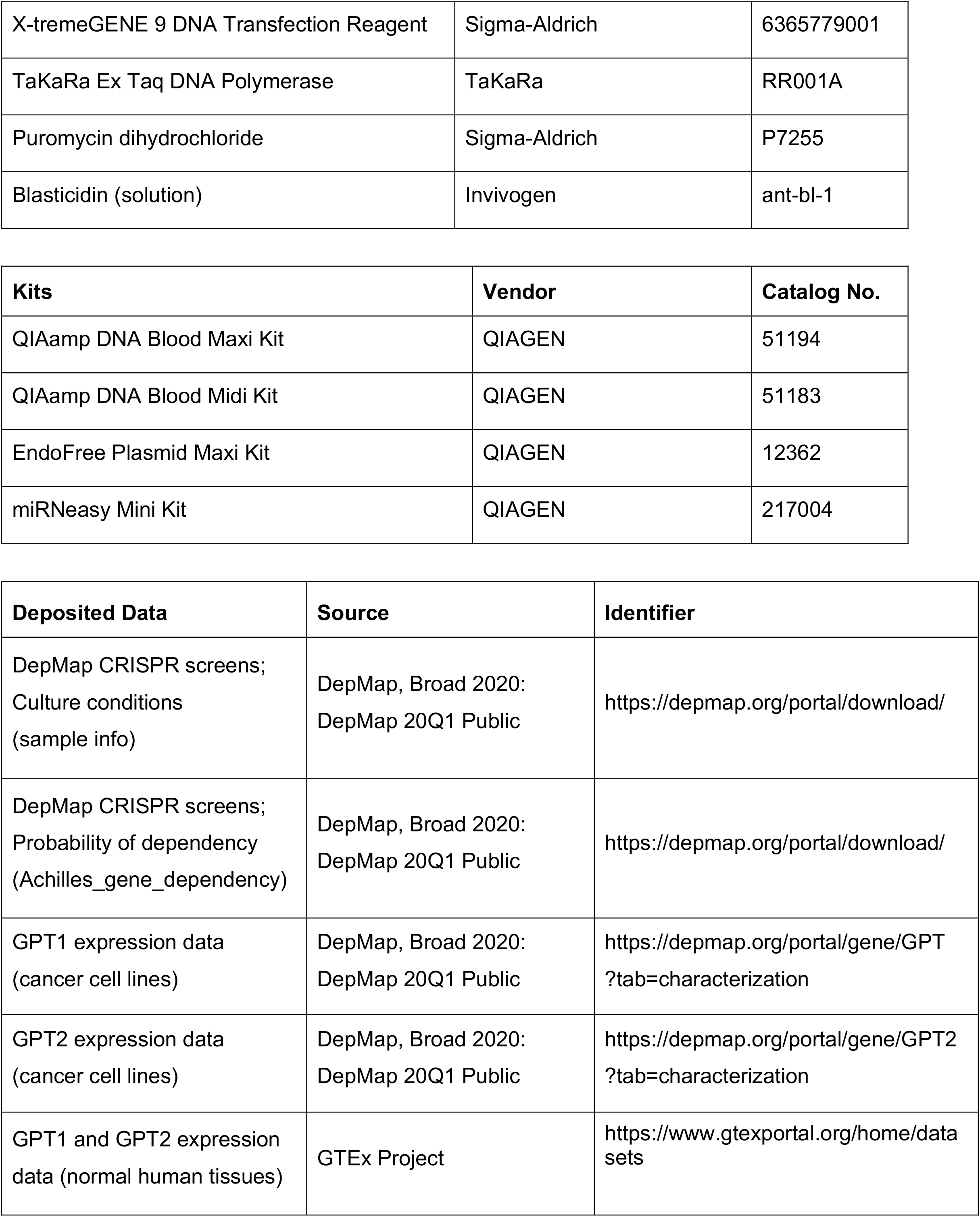
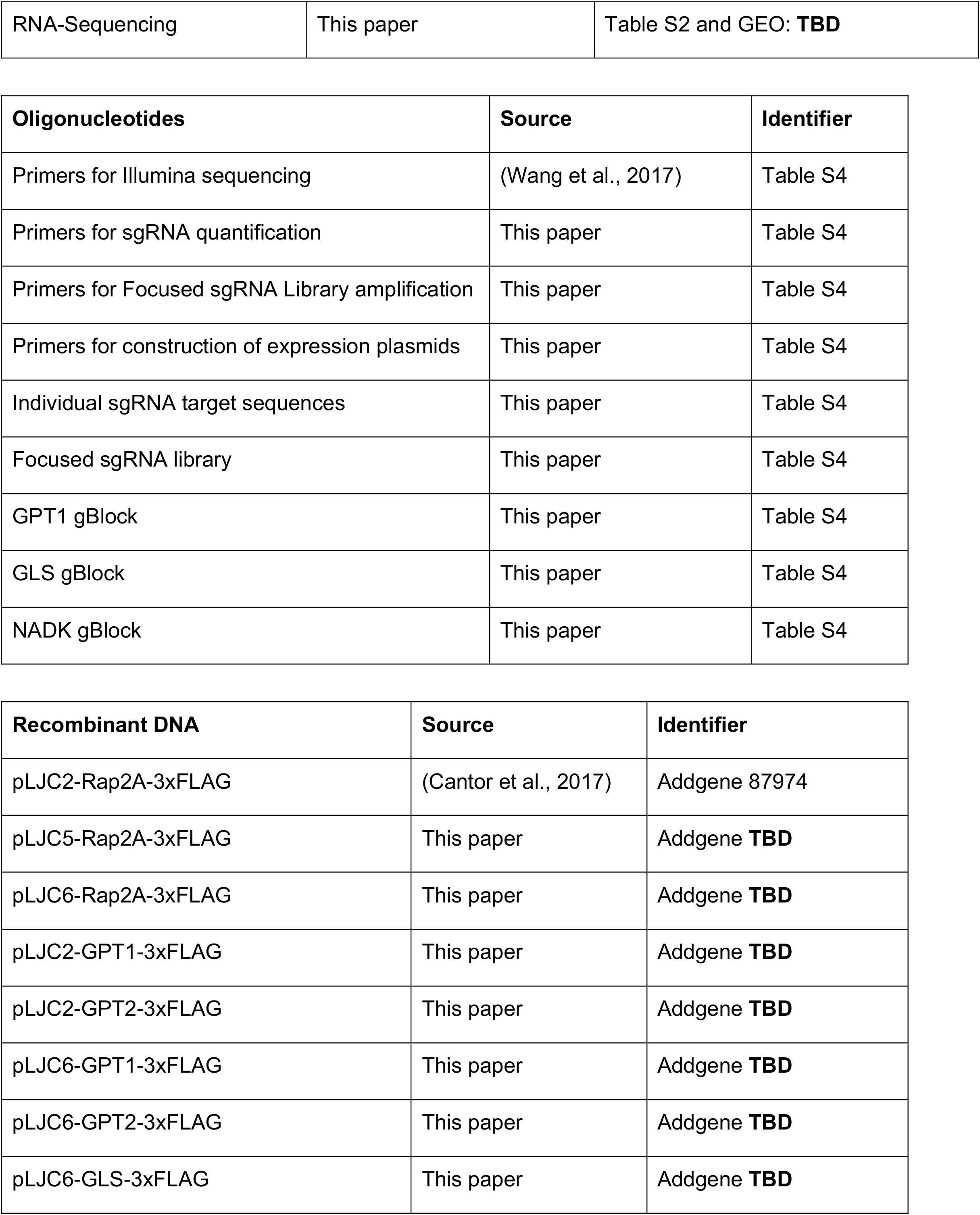
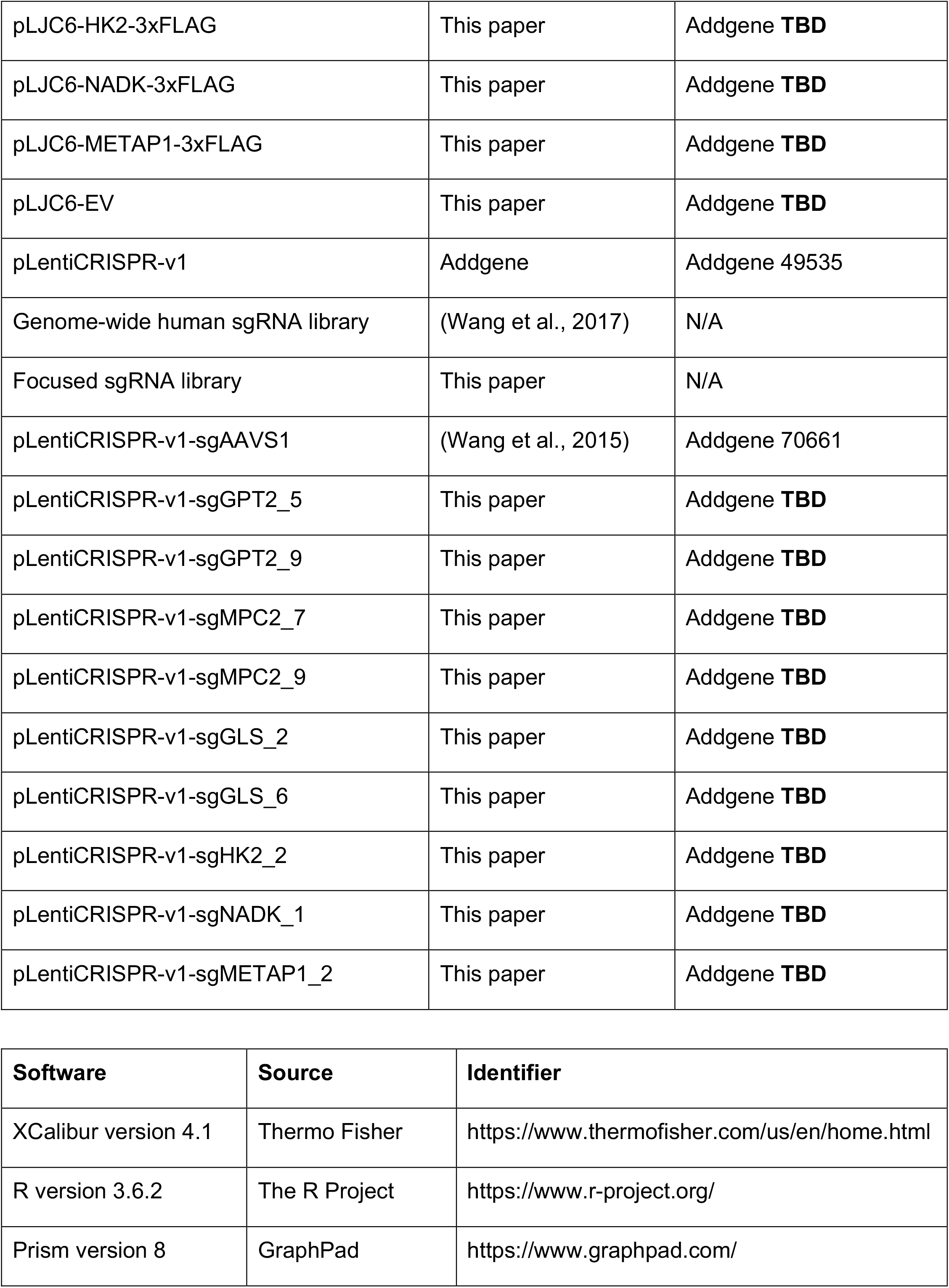
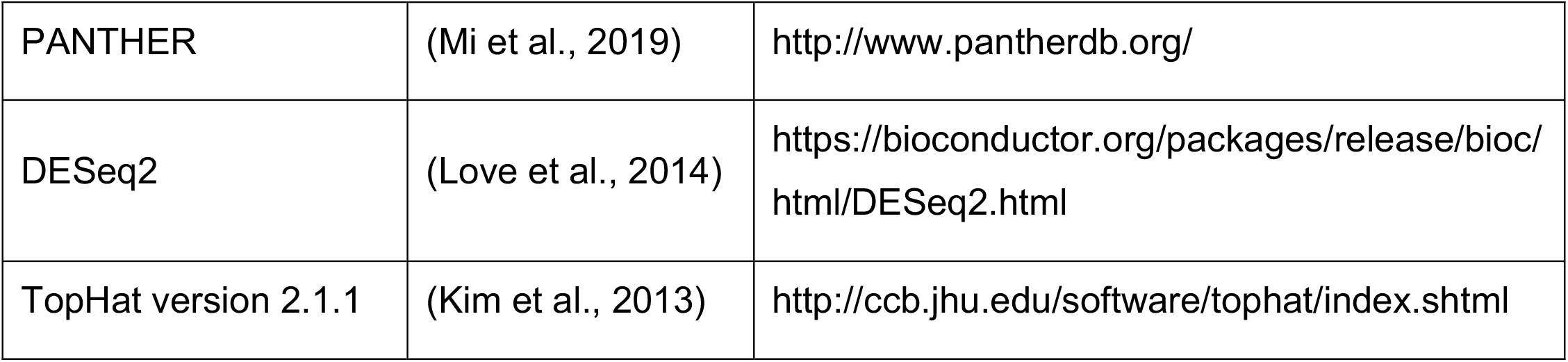

| Target | Vendor | Catalog No. |
| --- | --- | --- |
| Mouse monoclonal anti-GAPDH (0411) | Santa Cruz Biotechnology | sc-47724 |
| Rabbit polyclonal anti-RAPTOR | EMD Millipore | 09-217 |
| Rabbit monoclonal anti-GLS (6H5L15) | Thermo Fisher | 701965 |
| Mouse monoclonal anti-GPT (E-3) | Santa Cruz Biotechnology | sc-374501 |
| Mouse monoclonal anti-GPT2 (G-7) | Santa Cruz Biotechnology | sc-398383 |
| Rabbit monoclonal anti-MPC2 (D417G) | Cell Signaling Technology | 46141 |
| Rabbit monoclonal anti-HK2 (C64G5) | Cell Signaling Technology | 2867 |
| Rabbit polyclonal anti-NADK (55948) | Cell Signaling Technology | 55948 |
| Mouse monoclonal anti-METAP1 (A-2) | Santa Cruz Biotechnology | sc-514653 |
| Horse anti-Mouse IgG HRP | Cell Signaling Technology | 7076 |
| Goat anti-Rabbit IgG HRP | Cell Signaling Technology | 7074 |

| Bacterial strains | Vendor | Catalog No. |
| --- | --- | --- |
| Endura ElectroCompetent Cells | Lucigen | 60242 |
| XL10-Gold Ultracompetent Cells | Agilent | 200315 |

| Chemicals and Reagents | Vendor | Catalog No. |
| --- | --- | --- |
| 3X FLAG Peptide | Sigma-Aldrich | F4799 |
| Anti-FLAG M2 Affinity Gel | Sigma-Aldrich | A2220 |
| Mixture of amino acid standards for metabolomics | Cambridge Isotope Laboratories | MSK-A2-1.2 |
| Defined metabolite components of HPLM | Multiple | See Table S5 |
| Sodium Pyruvate | Sigma-Aldrich | P2256 |
| L-Glutamic acid | Sigma-Aldrich | G1251 |
| α-Ketoglutaric acid disodium salt dihydrate | Sigma-Aldrich | 75892 |
| Pyridoxal 5'-phosphate hydrate | Sigma-Aldrich | P9255 |
| D-glucose (U-$^{13}\text{C}_6$, 99%) | Cambridge Isotope Laboratories | CLM-1396 |
| L-Alanine ($^{13}\text{C}_3$, 99%) | Cambridge Isotope Laboratories | CLM-2184 |
| RPMI 1640 100X Vitamins | Sigma-Aldrich | R7256 |
| L-Alanine | Sigma-Aldrich | A7627 |
| Dimethyl α-ketoglutarate | Sigma-Aldrich | 349631 |
| Dimethyl malate | Sigma-Aldrich | 374318 |
| Dimethyl succinate | Sigma-Aldrich | W239607 |
| RPMI 1640, no glucose | Thermo Fisher | 11879020 |
| RPMI 1640 | Thermo Fisher | 11875093 |
| DMEM, no glucose | Thermo Fisher | 11966025 |
| DMEM, high glucose, GlutaMAX | Thermo Fisher | 10566024 |
| Glucose | Thermo Fisher | 15023021 |
| Penicillin-Streptomycin | Thermo Fisher | 15140122 |
| Fetal Bovine Serum (FBS), Heat Inactivated | Thermo Fisher | 16140071 |
| X-tremeGENE 9 DNA Transfection Reagent | Sigma-Aldrich | 6365779001 |
| TaKaRa Ex Taq DNA Polymerase | TaKaRa | RR001A |
| Puromycin dihydrochloride | Sigma-Aldrich | P7255 |
| Blasticidin (solution) | Invivogen | ant-bl-1 |

| **Kits** | **Vendor** | **Catalog No.** |
| --- | --- | --- |
| QIAamp DNA Blood Maxi Kit | QIAGEN | 51194 |
| QIAamp DNA Blood Midi Kit | QIAGEN | 51183 |
| EndoFree Plasmid Maxi Kit | QIAGEN | 12362 |
| miRNeasy Mini Kit | QIAGEN | 217004 |

| **Deposited Data** | **Source** | **Identifier** |
| --- | --- | --- |
| DepMap CRISPR screens;<br>Culture conditions<br>(sample info) | DepMap, Broad 2020:<br>DepMap 20Q1 Public | https://depmap.org/portal/download/ |
| DepMap CRISPR screens;<br>Probability of dependency<br>(Achilles_gene_dependency) | DepMap, Broad 2020:<br>DepMap 20Q1 Public | https://depmap.org/portal/download/ |
| GPT1 expression data<br>(cancer cell lines) | DepMap, Broad 2020:<br>DepMap 20Q1 Public | https://depmap.org/portal/gene/GPT<br>?tab=characterization |
| GPT2 expression data<br>(cancer cell lines) | DepMap, Broad 2020:<br>DepMap 20Q1 Public | https://depmap.org/portal/gene/GPT2<br>?tab=characterization |
| GPT1 and GPT2 expression<br>data (normal human tissues) | GTEx Project | https://www.gtexportal.org/home/data-sets |
| RNA-Sequencing | This paper | Table S2 and GEO: **TBD** |

| **Oligonucleotides** | **Source** | **Identifier** |
| --- | --- | --- |
| Primers for Illumina sequencing | (Wang et al., 2017) | Table S4 |
| Primers for sgRNA quantification | This paper | Table S4 |
| Primers for Focused sgRNA Library amplification | This paper | Table S4 |
| Primers for construction of expression plasmids | This paper | Table S4 |
| Individual sgRNA target sequences | This paper | Table S4 |
| Focused sgRNA library | This paper | Table S4 |
| GPT1 gBlock | This paper | Table S4 |
| GLS gBlock | This paper | Table S4 |
| NADK gBlock | This paper | Table S4 |

| **Recombinant DNA** | **Source** | **Identifier** |
| --- | --- | --- |
| pLJC2-Rap2A-3xFLAG | (Cantor et al., 2017) | Addgene 87974 |
| pLJC5-Rap2A-3xFLAG | This paper | Addgene **TBD** |
| pLJC6-Rap2A-3xFLAG | This paper | Addgene **TBD** |
| pLJC2-GPT1-3xFLAG | This paper | Addgene **TBD** |
| pLJC2-GPT2-3xFLAG | This paper | Addgene **TBD** |
| pLJC6-GPT1-3xFLAG | This paper | Addgene **TBD** |
| pLJC6-GPT2-3xFLAG | This paper | Addgene **TBD** |
| pLJC6-GLS-3xFLAG | This paper | Addgene **TBD** |
| pLJC6-HK2-3xFLAG | This paper | Addgene **TBD** |
| pLJC6-NADK-3xFLAG | This paper | Addgene **TBD** |
| pLJC6-METAP1-3xFLAG | This paper | Addgene **TBD** |
| pLJC6-EV | This paper | Addgene **TBD** |
| pLentiCRISPR-v1 | Addgene | Addgene 49535 |
| Genome-wide human sgRNA library | (Wang et al., 2017) | N/A |
| Focused sgRNA library | This paper | N/A |
| pLentiCRISPR-v1-sgAAVS1 | (Wang et al., 2015) | Addgene 70661 |
| pLentiCRISPR-v1-sgGPT2_5 | This paper | Addgene **TBD** |
| pLentiCRISPR-v1-sgGPT2_9 | This paper | Addgene **TBD** |
| pLentiCRISPR-v1-sgMPC2_7 | This paper | Addgene **TBD** |
| pLentiCRISPR-v1-sgMPC2_9 | This paper | Addgene **TBD** |
| pLentiCRISPR-v1-sgGLS_2 | This paper | Addgene **TBD** |
| pLentiCRISPR-v1-sgGLS_6 | This paper | Addgene **TBD** |
| pLentiCRISPR-v1-sgHK2_2 | This paper | Addgene **TBD** |
| pLentiCRISPR-v1-sgNADK_1 | This paper | Addgene **TBD** |
| pLentiCRISPR-v1-sgMETAP1_2 | This paper | Addgene **TBD** |

| **Software** | **Source** | **Identifier** |
| --- | --- | --- |
| XCalibur version 4.1 | Thermo Fisher | https://www.thermofisher.com/us/en/home.html |
| R version 3.6.2 | The R Project | https://www.r-project.org/ |
| Prism version 8 | GraphPad | https://www.graphpad.com/ |
| PANTHER | (Mi et al., 2019) | http://www.pantherdb.org/ |
| DESeq2 | (Love et al., 2014) | https://bioconductor.org/packages/release/bioc/html/DESeq2.html |
| TopHat version 2.1.1 | (Kim et al., 2013) | http://ccb.jhu.edu/software/tophat/index.shtml |

#### Cell lines

The following cell lines were kindly provided by: K562 and NOMO1, Dr. James Griffin (Dana Farber Cancer Institute); MOLM-13, the Cancer Cell Line Encyclopedia (Broad Institute); and SUDHL4, Dr. Margaret Shipp (Dana Farber Cancer Institute). Cells were verified to be free of mycoplasma contamination (Freshney, 2010) and cell line identities were authenticated by STR profiling.

### Methods

#### Cell culture

The following culture media were used in this study (each contained 0.5% penicillin-streptomycin):

(1) RPMI^+S^: RPMI 1640, no glucose (Thermo Fisher) with 5 mM glucose and 10% FBS.

(2) RPMI^+dS^: RPMI 1640, no glucose (Thermo Fisher) with 5 mM glucose and 10% dialyzed FBS.

(3) RPMI11^+2S^: RPMI 1640 (Thermo Fisher) with 20% FBS.

(4) DMEM^+S^: DMEM, no glucose (Thermo Fisher) with 5 mM glucose and 10% FBS.

(5) DMEM^+dS^: DMEM, no glucose (Thermo Fisher) with 5 mM glucose and 10% dialyzed FBS.

(6) DMEM25^+S^: DMEM, high glucose, GlutaMAX (Thermo Fisher) with 10% FBS.

(7) DMEM25^+2S^: DMEM, high glucose, GlutaMAX (Thermo Fisher) with 20% FBS.

(8) HPLM^+dS^: HPLM (See Table S5) with 10% dialyzed FBS and using RPMI 1640 100X Vitamins (Sigma-Aldrich R7256). Relative to the initially reported formulation (Cantor et al., 2017), HPLM was prepared with four additional components: α-KG, *O*-Acetylcarnitine, Malate, and Uridine.

(9) mHPLM^+dS^: minimal HPLM (See Table S5) with 10% dialyzed FBS and using RPMI 1640 100X Vitamins (Sigma-Aldrich R7256).

By using SnakeSkin tubing (Thermo Fisher PI88244), FBS was dialyzed as previously described (Cantor et al., 2017). Prior to use, all media were sterile filtered using bottle-top vacuum filters with cellulose acetate membrane, pore size 0.22 μm (Corning 430626). All cells were maintained at 37°C, atmospheric oxygen, and 5% CO_2_.

#### Genome-wide CRISPR screens

For genome-wide screens in K562 cells, the human sgRNA library described in (Wang et al., 2017) was used. To achieve at least 1000-fold coverage of the library following antibiotic selection, 350 million K562 cells were seeded at a density of 2.5 × 10^6^ cells/mL in 6-well plates containing 2 mL of RPMI^+S^ containing 8 μg/mL polybrene and the pLentiCRISPR-v1 library virus. Spin infection was carried out by centrifugation at 2,200 RPM for 45 min at 37°C. After 18 hr incubation, the cells were pelleted to remove virus and then re-seeded into fresh RPMI^+S^ for 24 hr. Cells were then pelleted and re-seeded to a density of 150,000 cells/mL in RPMI^+S^ containing 2 μg/mL puromycin (Sigma-Aldrich) for 72 hr. Following selection, an initial pool of 100 million cells was pelleted and frozen, and pools of 216 million cells were used to collectively seed each of twelve total 225 cm^2^ rectangular canted neck cell culture flasks (Corning 431082) to a density of 150,000 cells/mL in 120 mL of either HPLM^+dS^ or RPMI^+dS^, respectively. Cells were passaged every 48 hr and population doublings were tracked by cell density measurements using a Coulter Counter (Beckman Z2) with a diameter setting of 8-30 μm. After 13 population doublings, a pool of 100 million cells from each screen was harvested for genomic DNA (gDNA) extraction using the QIAamp DNA Blood Maxi Kit (QIAGEN).

Using Ex Taq DNA Polymerase (Takara), sgRNA inserts from each initial and final pool were PCR-amplified from 290 μg of gDNA to achieve ∼400-fold coverage of the library. The resulting PCR products were purified and sequenced on a HiSeq 2500 (Illumina) (Primer sequences are annotated in Table S6) to quantify sgRNA abundances in each sample.

#### RNA sequencing

K562 cells were pelleted and then seeded at a density of 200,000 cells/mL in 6 cm culture dishes containing 6 mL fresh medium. After 24 hr incubation, RNA was harvested using the miRNeasy Mini Kit (QIAGEN) according to manufacturer instructions. 700 ng total RNA was used to generate mRNA libraries using the TruSeq Stranded mRNA Library Preparation Kit (Illumina 20020594). Libraries prepared from each sample were quantified using the KAPA Library Quantification Kit (Roche KK4824) and then pooled at equimolar ratios. Following denaturation, 40 bp single-end reads were generated on a HiSeq 2500 (Illumina). Base calls were performed by the instrument control software and further processed using the Offline Base Caller version 1.9.4 (Illumina), and quality control analysis was performed using the FastQC program (Babraham Bioinformatics).

Reads were aligned to the human genome (GRCh37) with Ensembl annotation v. 75 using Tophat version 2.1.1 (parameters ‘--no-novel-juncs’ and ‘--segment-length’ of 20) (Kim et al., 2013). Across all samples, the overall mapping rate was 91.6% (average) and 79.1% (average) of the reads aligned uniquely. Based on two biological replicates for each condition, read counts were quantified at the gene level and normalized using the geometric means method implemented in the DESeq2 package (Love et al., 2014).

#### Focused sgRNA library construction

To design the focused sgRNA library, subsets of 212 HPLM- and 182 RPMI-essential hit genes that collectively span a number of manually curated biological processes were first selected from genome-level screen results. An additional 257 genes related to these hits through either a shared pathway, gene family, or encoded protein complex were then selected as well. The focused sgRNA library contained 16,585 constructs targeting 651 total protein-coding genes (up to 25 sgRNAs per gene) and 325 non-targeting sgRNAs. An oligonucleotide pool for the library was synthesized (Agilent), PCR-amplified according to manufacturer instructions using the primers JC842/JC843 to incorporate overhangs compatible for Gibson Assembly (New England Biolabs), and cloned into *BsmBI*-digested pLentiCRISPR-v1. Reaction products were transformed into *E. coli* Endura electrocompetent cells (Lucigen), plated onto prewarmed LB medium/agar containing 100 μg/mL ampicillin in a 245 mm square bioassay dish (Corning 431111), and incubated for 18 hr at 30°C, yielding ∼10^7^ individual transformants – equivalent to ∼500-fold coverage of the theoretical library diversity. Colonies were scraped and pooled in LB medium, and plasmid DNA was then extracted using the EndoFree Maxi Kit (QIAGEN).

#### Secondary CRISPR screens

The secondary screening procedure was similar to that used for the genome-wide screens with minor modifications.

1. To achieve at least 1000-fold coverage of the library following antibiotic selection in each cell line, 60 million cells were transduced with virus
2. For selection of the other transduced cell lines, puromycin was used at the following concentrations: NOMO1 (0.5 μg/mL), MOLM13 (1 μg/mL), and SUDHL4 (0.5 μg/mL).
3. To initiate and maintain each screen, 18 million cells were seeded in single flasks to the same density (150,000 cells/mL) and in the same volume (120 mL) of culture medium
4. To harvest gDNA from each pool, 10 million cells were extracted using the QIAamp DNA Blood Midi Kit (QIAGEN)
5. For each extraction, sgRNA inserts were PCR-amplified from 24 μg of gDNA

#### Plasmid construction

All oligonucleotides used in this study are described in Table S4.

##### Construction of lentiviral plasmids pLJC5-Rap2A-3xFLAG and pLJC6-Rap2A-3xFLAG

The human ubiquitin C (UbC) promoter was amplified from the pLenti6/UbC/V5-DEST Gateway vector (Thermo Fisher V49910) using the primers JC419/JC420, digested with *ClaI*-*AgeI*, and cloned into pLJC2-Rap2A-3xFLAG to generate pLJC5-Rap2A-3xFLAG. The blasticidin resistance cassette was amplified from the pMXs-IRES-blasticidin retroviral vector (Cell Biolabs RTV-016) using the primers JC606/JC607, digested with *BamHI*-*KpnI*, and cloned into pLJC5-Rap2A-3xFLAG to generate pLJC6-Rap2A-3xFLAG.

##### Construction of gene knockout plasmids

For each of the following genes, sense and antisense oligonucleotides were annealed and then cloned into *BsmBI*-digested pLentiCRISPR-v1: *GPT2, MPC2, GLS, HK2, NADK*, and *METAP1*.

##### Construction of expression plasmids

The *GPT2* gene was amplified using the primers GPT2-F/GPT2-R, digested with *PacI*-*NotI*, and cloned into pLJC2-Rap2A-3xFLAG to generate pLJC2-GPT2-3xFLAG. GPT2-F was designed to remove a *GPT2*-internal *NotI* site and to reduce the GC content at the 5’ terminus of the gene. Plasmid pLJC6-GPT2-3xFLAG contains a sgGPT2_5-resistant *GPT2* cDNA and was generated using a 2-step protocol based on overlap extension PCR methodology. In the first step, two fragments were amplified from pLJC2-GPT2-3xFLAG using the following primer pairs: LJC-F/GPT2_5-R and GPT2_5-F/LJC-R. In the second step, the two fragments were pooled in a second PCR containing the primers LJC-F/LJC-R, then digested with *PacI*-*NotI*, and cloned into pLJC6-Rap2A-3xFLAG. The same 2-step protocol was used to generate plasmids pLJC6-HK2-3xFLAG and pLJC6-METAP1-3xFLAG, which contain a sgHK2_2-resistant-*HK2* cDNA and a sgMETAP1_2 *METAP* cDNA, respectively. For pLJC6-HK2-3xFLAG, the following internal primers were used: HK2_2-F and HK2_2-R. For pLJC6-METAP1-3xFLAG, the following internal primers were used: METAP1_2-F and METAP1_2-R.

The GPT1 gene was amplified from a codon-optimized gBlock Gene Fragment (IDT) using the primers GPT1-F/GPT1-R, digested with *PacI*-*NotI*, cloned into pLJC2-Rap2A-3xFLAG to generate pLJC2-GPT1-3xFLAG, and then subcloned into pLJC6-Rap2A-3xFLAG to generate pLJC6-GPT1-3xFLAG as well. The GLS gene was amplified from a codon-optimized gBlock Gene Fragment (IDT) using the primers GLS-F/GLS-R, digested with *PacI*-*NotI*, and cloned into pLJC6-Rap2A-3xFLAG to generate pLJC6-GLS-3xFLAG, which contains a sgGLS_6-resistant *GLS* cDNA. The NADK gene was amplified from a codon-optimized gBlock Gene Fragment (IDT) using the primers NADK-F/NADK-R, digested with *PacI*-*NotI*, and cloned into pLJC6-Rap2A-3xFLAG to generate pLJC6-NADK-3xFLAG, which contains a sgNADK_1-resistant *NADK* cDNA.

To create an empty vector (EV) derivative of pLJC6, oligonucleotides JC1145 and JC1146 were annealed and cloned into pLJC6-Rap2A-3xFLAG at *PacI*-*NotI* to generate pLJC6-EV.

#### Lentivirus production

To produce lentivirus, HEK293T cells in DMEM25^+S^ were co-transfected with the VSV-G envelope plasmid, the Delta-VPR packaging plasmid, and the appropriate transfer plasmid (either a pLJC6 or pLentiCRISPR-v1 backbone) using X-tremeGENE 9 Transfection Reagent (Sigma-Aldrich). Culture medium was exchanged with DMEM25^+2S^ 16 hr after transfection, and the virus-containing supernatant was collected at 48 hr post-transfection, passed through a 0.45 μm filter to eliminate cells, and then stored at −80°C.

#### Cell line construction

##### Knockout cell lines

To establish knockout clonal cell lines, K562 cells were seeded at a density of 500,000 cells/mL in 6-well plates containing 2 mL RPMI^+S^, 8 μg/mL polybrene, and the appropriate pLentiCRISPR-v1 lentivirus. Spin infection was carried out by centrifugation at 2,200 RPM for 45 min at 37°C. After 16-18 hr incubation, the cells were pelleted to remove virus and then re-seeded into fresh RPMI^+S^ for 24 hr. Cells were then pelleted and re-seeded into fresh RPMI^+S^ containing puromycin (Sigma-Aldrich) for 72 hr and, following selection, were single-cell FACS-sorted into 96-well plates containing RPMI11^+2S^. After 1.5-2 weeks, cell clones with the desired knockouts were identified by immunoblotting. To control for infection with pLentiCRISPR-v1 virus, a control population of K562 cells was similarly selected following transduction with sgAAVS1-containing virus (Wang et al., 2015).

##### cDNA expression cell lines

To establish stable expression cell lines, K562 clonal cells were seeded at a density of 175,000 cells/mL in 6-well plates containing 2 mL of RPMI^+S^, 8 μg/mL polybrene, and the appropriate pLJC6 lentivirus. Spin infection and initial media exchange were each performed identically to those for knockout cell lines. Cells were then pelleted and re-seeded into fresh RPMI^+S^ containing blasticidin (Invivogen) for 72 hr. Stable expression of cDNA was confirmed by immunoblotting.

#### Cell lysis for immunoblotting

Cells were centrifuged at 250 *g* for 5 min, resuspended in 1 mL ice-cold PBS, and then centrifuged again at 250 *g* for 5 min at 4°C. Cells were then immediately lysed with ice-cold lysis buffer (40 mM Tris-HCl pH 7.4, 1% Triton X-100, 100 mM NaCl, 5 mM MgCl_2_, 1 tablet of EDTA-free protease inhibitor (Roche 11580800; per 25 mL buffer), 1 tablet of PhosStop phosphatase inhibitor (Roche 04906845001; per 10 mL buffer). The cell lysates were cleared by centrifugation at 21130 *g* for 10 min at 4°C and then quantified for protein concentration using an albumin standard (Thermo Fisher 23209) and Bradford reagent (Bio-Rad 5000006). Cell lysate samples were normalized for protein content, denatured upon the addition of 5X sample buffer (Thermo Fisher 39000), resolved by 12% SDS-PAGE, and transferred to a polyvinyl difluoride membrane (Millipore IPVH07850). Membranes were blocked with 5% nonfat dry milk in TBST for 1 hr at room temperature, and then incubated with primary antibodies in 5% nonfat dry milk in TBST overnight at 4°C. Primary antibodies to the following proteins were used at the indicated dilutions: GAPDH (1:1000); RAPTOR (1:1000); GPT1 (1:100); GPT2 (1:100); MPC2 (1:100); GLS (1:100); HK2 (1:1000); NADK (1:300); and METAP1 (1:200). Membranes were washed with TBST three times for 5 min each, and then incubated with species-specific HRP-conjugated secondary antibody (1:3000) in 5% nonfat dry milk for 1 hr at room temperature. Membranes were washed again with TBST three times for 5 min each, and then visualized with chemiluminescent substrate (Thermo Fisher) on a LICOR Odyssey FC.

#### Short-term proliferation assays

Following at least two passages in RPMI^+S^, cells were pelleted and resuspended to a density of 1 million cells/mL in RPMI^+S^. From each resuspension, 80,000 total cells were seeded in each of three replicate wells containing 4 mL of the appropriate culture medium in 6-well plates. Following 96 hr incubation, cell density measurements were recorded using a Coulter Counter (Beckman Z2) with a diameter setting of 8-30 μm. Stock solutions of the following components were prepared relative to working concentrations in HPLM: L-Alanine (500X), αKG (1000X), 2-hydroxybutyrate, 3-hydroxybutyrate, malonate (250X), citrate (250X), malate (1000X), succinate (1000X), acetate (500X), lactate (100X), and pyruvate (250X). In addition, stock solutions of dimethyl αKG, dimethyl malate, and dimethyl succinate were each prepared at 100 mM in water.

#### Metabolite Profiling and Quantification of Metabolite Abundance

LC-MS analyses were performed on a QExactive HF benchtop orbitrap mass spectrometer equipped with an Ion Max API source and HESI II probe, which was coupled to a Vanquish Horizon UPLC system (Thermo Fisher). External mass calibration was performed using positive and negative polarity standard calibration mixtures every 7 days. Acetonitrile was hypergrade for LC-MS (Millipore Sigma) and all other solvents were Optima LC-MS grade (Thermo Fisher).

##### Cells

Following at least two passages in RPMI^+S^, cells were pelleted, resuspended in fresh medium of interest, and then seeded in a volume of 4 mL per well at a density of 125,000 cells/mL in 6-well plates. For labeling experiments, the procedure was identical except that RPMI^+dS^ containing either 5 mM [U-^13^C]-glucose or 430 μM ^13^C_3_-alanine was used. After 24 hr incubation, a 500 μL aliquot was used to measure cell number and volume via Coulter Counter (Beckman Z2) with a diameter setting of 8-30 μm, and the remaining cells were centrifuged at 250 *g* for 5 min, resuspended in 1 mL ice-cold 0.9% sterile NaCl (Growcells MSDW1000), and again centrifuged at 250 *g* for 5 min at 4°C. Metabolites were extracted in 1 mL ice-cold 80% methanol containing 500 nM internal amino acid standards (Cambridge Isotope Laboratories). Following a 10 min vortex and centrifugation for 3 min at 21130 *g* for 10 min at 4°C, samples were dried under nitrogen gas. Dried samples were stored at −80°C and then resuspended in 100 μL water. Following a 10 min vortex and centrifugation at 21130 *g* for 10 min at 4°C, 2.5 μL from each cell sample was injected onto a ZIC-pHILIC 2.1 x 150 mm analytical column equipped with a 2.1 x 20 mm guard column (both 5 μm particle size, Millipore Sigma). Buffer A was 20 mM ammonium carbonate, 40 mM ammonium hydroxide; buffer B was acetonitrile. The chromatographic gradient was run at a flow rate of 0.15 mL/min as follows: 0-20 min: linear gradient from 80% to 20% B; 20-20.5 min: linear gradient from 20% to 80% B; 20.5-28 min: hold at 80% B.

The mass spectrometer was operated in full scan, polarity-switching mode with the spray voltage set to 3.0 kV, the heated capillary held at 275°C, and the HESI probe held at 350°C. The sheath gas flow rate was set to 40 units, the auxiliary gas flow was set to 15 units, and the sweep gas flow was set to 1 unit. The MS data acquisition in positive mode was performed in a range of 50-750 m/z, with the resolution set to 120,000, the AGC target at 10^6^, and the maximum integration time at 20 msec. The settings in negative mode were the same except that the range was instead 70-1000 m/z.

##### Media

To extract metabolites from cell culture media, samples were diluted 1:40 in a solution of 50:30:20 methanol:acetonitrile:water containing 500 nM internal amino acid standards (Cambridge Isotope Laboratories). Following a 10 min vortex and centrifugation at 21130 *g* for 5 min at 4°C, 2.5 μL of each sample was injected for LC-MS analysis as described above for profiling cell samples.

##### GPT activity assay

For detection of αKG generated in GPT activity assays, reaction mixtures were extracted (See GPT Activity Assay) and 5 μL was injected onto a ZIC-pHILIC 2.1 x 150 mm analytical column equipped with a 2.1 x 20 mm guard column (both 5 μm particle size, Millipore Sigma). Buffer A was 20 mM ammonium carbonate, 40 mM ammonium hydroxide; buffer B was acetonitrile. The chromatographic gradient was run at a flow rate of 0.15 mL/min as follows: 0-10 min: linear gradient from 80% to 20% B; 10-10.5 min: linear gradient from 20% to 80% B; 10.5-16.5 min: hold at 80% B. The mass spectrometer was operated in full scan, polarity-switching mode with the spray voltage set to 3.5 kV (positive mode) and 2.5 kV (negative mode), the heated capillary held at 275°C, and the HESI probe held at 350°C. The sheath gas flow rate was set to 40 units, the auxiliary gas flow was set to 10 units, and the sweep gas flow was set to 1 unit. The MS data acquisition in negative mode was performed in a range of 70-1050 m/z, with the resolution set to 120,000, the AGC target at 10^6^, and the maximum integration time at 200 msec.

##### Highly Targeted Metabolomics

For the highly targeted analyses of acetyl-CoA and pyruvate in whole-cell samples without nutrient labeling, the instrument was run as described (See **Cells**), but with additional tSIM (targeted selected ion monitoring) scans in negative ionization mode. The tSIM settings were as follows: resolution set to 120,000, an AGC target of 10^5^, and a maximum integration time of 200 msec. The target masses were 808.1185 (corresponding to acetyl-CoA) and 87.0088 (corresponding to pyruvate). The isolation window around each target mass was set to 1.0 m/z.

For αKG and pyruvate in culture media samples, all settings as described for the tSIM scan used for cell samples were identical, except that the target mass 145.0142 (corresponding to αKG) was added. For pyruvate and M+3-pyruvate in cell samples with nutrient labeling, all settings were again identical, except that the target mass 90.01887 (corresponding to M+3-pyruvate) was added.

For αKG in GPT activity assay samples, all settings as described for the tSIM scan used for αKG in media were identical except that the maximum integration time was 400 msec. For pyruvate in the same samples, all settings as described for the tSIM scan used for pyruvate were identical. Finally, for L-Phenylalanine (^13^C_9_, 99%; ^15^N, 99%) in the same samples, all settings as described for the tSIM scan used for whole-cell metabolites were identical except that the scan was run in positive ionization mode, and the target mass 176.1135 was added.

##### Identification and Quantification

Metabolite identification and quantification were performed with XCalibur version 4.1 (Thermo Fisher) using a 10-ppm mass accuracy window and 0.5 min retention time window. To confirm metabolite identities and to enable quantification when desired, a manually constructed library of chemical standards was used. Standards were validated by LC-MS to confirm that they generated robust peaks at the expected *m/z* ratio, and stock solutions were stored in pooled format at −80°C at the following concentrations: 1 mM, 100 μM, 10 μM, and 0.1 μM. On the day of a given queue, each stock was diluted 1:10 in water containing 500 nM internal amino acid standards (Cambridge Isotope Laboratories), and then vortexed and centrifuged as described for biological samples (See **Cells** and **Media**). For those metabolites lacking a chemical standard, peak identification was restricted to high confidence peak assignments (Smith et al., 2005). See table S6.

Because metabolite extraction protocols differed by sample type, the internal standard concentrations in processed samples varied: chemical standards (450 nM), media samples (487.5 nM), and cell samples (5 μM). Therefore, the raw peak areas of internal standards within each sample of a given batch were first normalized to account for these differences. Metabolite quantification was then performed as described elsewhere (Cantor et al., 2017).

#### Expression and immunoprecipitation of recombinant proteins

For isolation of recombinant GPT proteins, 4 million HEK293T cells were plated in 15 cm culture dishes containing DMEM25^+S^. After 24 hr incubation, cells were transfected with 15 μg of pLJC2-GPT1-3xFLAG or pLJC2-GPT2-3xFLAG as described elsewhere (Cantor et al., 2017). After an additional 48 hr incubation, cells were rinsed once with ice-cold PBS and then immediately lysed in ice-cold lysis buffer (See **Cell lysis for immunoblotting**). The cell lysates were cleared by centrifugation at 21130 *g* for 10 min at 4°C. For anti-FLAG immunoprecipitation, the FLAG-M2 affinity gel (Sigma-Aldrich) was washed three times in lysis buffer, and then 400 μL of a 50:50 affinity gel slurry was added to a pool of clarified lysates collected from five individual 15 cm culture dishes, and incubated with rotation for 3 hr at 4°C. Following immunoprecipitation, the beads were washed twice in lysis buffer and then four times with lysis buffer containing 500 mM NaCl. Recombinant protein was then eluted in lysis buffer containing 500 μg/mL 3x-FLAG peptide (Sigma-Aldrich) for 1 hr with rotation at 4°C. The eluent was isolated by centrifugation at 100 *g* for 4 min at 4°C (Bio-Rad 732-6204), buffer exchanged (Amicon Ultra 30 kDa MWCO UFC503024) against 20 volumes of storage buffer (40 mM Tris-HCl pH 7.5, 100 mM NaCl, 2 mM dTT, 100 μM pyridoxal 5’-phosphate (PLP)), mixed with glycerol (final concentration 15% v/v), snap-frozen with liquid nitrogen, and stored at −80°C.

Protein samples were quantified using an albumin standard (Thermo Fisher 23209) and Bradford reagent (Bio-Rad 5000006). Purified proteins were normalized for protein content, denatured upon the addition of 5X sample buffer (Thermo Fisher 39000), and resolved by 12% SDS-PAGE. Apparent molecular weights via immunoblotting were comparable to those expected, but upon loading 600-fold more purified protein, those via Coomassie staining were lower than expected, likely reflective of differences in buffer conditions in the samples.

#### GPT Activity Assay

To determine kinetic constants for the conversion of pyruvate and L-Glutamate (L-Glu) to L-Alanine and αKG catalyzed by each GPT, we developed an in vitro GPT activity assay. To estimate kinetic parameters for pyruvate, reactions of purified GPT (2-4 nM enzyme) with fixed L-Glu (2.5 mM) and varying concentrations of pyruvate were carried out at 37°C in 40 mM Tris-HCl pH 7.5, 5 mM MgCl_2_, 5 mM Na_2_HPO_4_, 2 mM dTT, 500 μM NaCl, 150 μM PLP, and 100 μM EDTA in a total volume of 100 μL. After 30 sec incubation, a 35 μL aliquot of the reaction was removed and immediately added to 65 μL ice-cold 50:30:20 methanol:acetonitrile:water containing 500 nM internal amino acid standards (Cambridge Isotope Laboratories) for metabolite extraction. Samples were then vortexed for 10 min and centrifuged at 21130 *g* for 1 min at 4°C. To estimate kinetic parameters for L-Glu, the assay extraction procedure was similar with minor modifications. Reactions of purified GPT (20-40 nM) with fixed pyruvate (1 mM) and varying concentrations of L-Glu were carried out in the same conditions, except incubation times were either 1 min (GPT1) or 5 min (GPT2).

Concentrations of αKG generated in each reaction were evaluated by LC-MS analysis of extracted samples (See **Metabolite Profiling and Quantification of Metabolite Abundance**).

Using peak areas of an αKG standard normalized by those of L-Phenylalanine (^13^C_9_, 99%; ^15^N, 99%) identically prepared in the same extraction solution, we constructed standard curves fit to linear equations to ensure that αKG concentrations in the reaction samples did not exceed ∼10% of the initial substrate concentrations. For reactions with fixed L-Glu, standard curves consisted of points at the following concentrations: 167 nM, 500 nM, 1.5 μM, and 4.5 μM; and for those with fixed pyruvate, they were instead at: 4.5 μM, 13.5 μM, 40.5 μM, and 121.5 μM.

Of note, given both the 10% turnover threshold and αKG detection limit from the methods used, we could not measure meaningful αKG concentrations from reactions containing pyruvate below the estimated *K*_M, PYR_ of GPT2. Stock solutions of L-Glu (Sigma-Aldrich) and PLP (Sigma-Aldrich) were prepared at 100 mM in 10 mM HCl and 200 mM HCl, respectively, and upon appropriate dilutions, had little effect on reaction pH.

### Statistical Analyses

#### Genome-wide CRISPR screens

Sequencing reads were aligned to the sgRNA library and only exact matches were allowed. sgRNAs with less than 50 counts in the initial dataset were removed from downstream analysis. Genes targeted by less than seven distinct sgRNAs following this filtering were also removed from further analysis. Abundances of all remaining sgRNAs were determined by adding a pseudocount of one and then normalizing by the total number of read counts for a given sample. Depletion scores were calculated as the log_2_ fold-change in abundance of each sgRNA between the initial and final populations. Gene scores were defined as the average log_2_ fold-change in depletion scores of all sgRNAs targeting the gene.

Screens in different conditions may introduce discrepancies in aggregate gene selection that affect the dynamic range of gene scores (Wang et al., 2019). Thus, to reduce potential bias in calculating differential scores on an assumption that such distributions are equivalent between screens, we scaled all gene scores based on the assumption that sets of nontargeting (NT) sgRNAs and core essential genes (CEGs) would exhibit the same selection across different screens. In brief, gene scores were scaled such that the medians of post-filtering NT sgRNAs (989) and reference CEGs (680) (Hart et al., 2017) were defined as 0 and −1, respectively.

For each gene, a differential score between screens was calculated and then standardized relative to the entire set of targeted genes to assess differential dependency.

#### Probability of Dependency in genome-wide screens

For each genome-wide screen, probabilities of dependency were calculated for all targeted genes (Dempster et al., 1977, 2019). In brief, the gene score dataset from each screen was treated as a mixture model comprised of two normal distributions. Densities were generated using a standard E-M optimization procedure initialized with the means and standard deviations from reference sets of 680 CEGs and 768 nonessential genes (Hart et al., 2014, 2017). The probability of dependency for a given gene was then calculated as the ratio of CEG density to the sum of the two densities at the gene score of interest. Given that standard deviations of the two distributions differ, their estimated densities converge to zero at different rates in extreme tail regions, which can cause erroneous inflation of estimated probabilities at large enough gene score values. Thus, genes with a score greater than or equal to zero were assigned a dependency probability of 0.

#### Genes involved in fundamental processes

The following KEGG gene sets were obtained from the Gene Set Enrichment Analysis (GSEA) database: aminoacyl tRNA biosynthesis, DNA replication, nucleotide excision repair, proteasome, ribosome, RNA polymerase, and spliceosome.

#### Receiver-operator analysis

From each genome-wide screen dataset, receiver-operator characteristic (ROC) curves were generated from relatively balanced reference sets of 680 CEG and 768 nonessential genes (Hart et al., 2014, 2017). Area under the ROC curve was used as the performance metric to assess how well each could discriminate for CEGs.

#### PANTHER pathway-enrichment analysis

To determine which biological processes were enriched among conditionally essential hit genes from genome-level screens, genes were queried using the PANTHER Overrepresentation Test with Gene Ontology Biological Processes as the annotation dataset (Mi et al., 2019). Significance was measured with Fisher’s Exact Test using a false discovery rate (FDR) cutoff of 0.05.

#### RNA sequencing

Differential expression was evaluated by FeatureCounts. Significance of differential expression between conditions was measured using negative binomial distribution as implemented in the DESeq2 package, with *P*-values multiple-test corrected to estimate FDRs using the Benjamini-Hochberg procedure.

#### Secondary CRISPR screens

Gene scores were calculated using a procedure similar to that for the genome-wide screens with minor modifications. From each initial reference set, sgRNAs with less than 100 counts were removed from downstream analysis. sgRNA depletion scores were similarly scaled, but with the post-filtering NT sgRNAs in each initial dataset and CEGs (83) targeted by the focused library.

To combine data from replicate secondary screens in the K562 cell line in each of HPLM^+dS^ and RPMI^+dS^, scaled sgRNA-level data from replicates were pooled and gene scores were then calculated as the average from all sgRNAs targeting the gene. For all genes, *P*-values to compare differential depletion distributions of respective targeting sgRNAs to those of NT sgRNAs were calculated using a two-tailed Welch’s *t*-test, and then multiple-test corrected to estimate FDRs using the Benjamini-Hochberg procedure. Significance of the *r* value that describes the relationship between conditional phenotypes from genome-scale and pooled secondary screens was determined from a correlation test performed in R.

#### Quality control for linear transformations of gene score datasets

To assess the separation of control set distributions from each screen, strictly standardized mean difference (SSMD) statistics were calculated using the sgRNA depletion scores from NT sgRNAs and CEGs (Zhang, 2007, 2008). For all CRISPR screens in this study, calculated SSMD values were < −2, indicating excellent separation.

#### Metabolite Profiling

To compare intracellular metabolite abundances, *P*-values were calculated using a two-tailed Welch’s *t*-test, and then multiple-test corrected to estimate FDRs using the Benjamini-Hochberg procedure. *P*-values to compare differences in fractional labeling were also calculated using a two-tailed Welch’s *t*-test.

#### Enzyme kinetics

To determine kinetic constants for pyruvate and L-Glu, plots of substrate concentration versus reaction velocity in GraphPad Prism were fit using the Michaelis-Menten equation.

*P*-values to compare relative growth were determined using a two-tailed Welch’s *t*-test. The exact value of *n* and the definition of center and precision measures are provided in associated figure legends. Bar graphs were prepared in GraphPad Prism 8; remaining plots and heatmaps were prepared in R. All instances of reported replicates refer to *n* biological replicates.

### Data and Materials Availability

Data resources can be found in Tables S1, S2, S3, and S6. RNA-Seq data from this study is available from the Gene Expression Omnibus (GEO: TBD). Plasmids generated in this study will be deposited at Addgene.

